# Bach2 is a potent repressor of Nrf2-mediated antioxidant enzyme expression in dopaminergic neurons

**DOI:** 10.1101/687590

**Authors:** Elisa A. Waxman

**Affiliations:** Department of Neurobiology, Medical Diagnostic Laboratories, LLC, Hamilton, NJ, USA

**Keywords:** Nrf2, Bach1, Bach2, antioxidants, oxidative stress, Parkinson’s disease, heme oxygenase-1, NQO1, transcription, glutamyl cysteine ligase

## Abstract

Nuclear factor erythroid 2–related factor 2 (Nrf2) is a transcriptional activator of antioxidant response elements (ARE), which function to increase expression of antioxidant enzymes. Recent works suggests Nrf2 activation is a promising protective mechanism against the progressive neurodegeneration in Parkinson’s disease (PD). While Nrf2 inducers show some promising results in animal models of PD, the response is limit in dopaminergic neurons with the protection mostly conferred by surrounding glia. The present study characterizes ARE transcriptional repressors Bach1 and Bach 2 (Broad complex– Tramtrack–Bric-a-brac and Cap n’ Collar homology, basic leucine zipper transcription factors 1 and 2) as a potential explanation for limited Nrf2 activity in these neurons. The current work identified Bach1 and Bach2 in dopaminergic neurons of the human substantia nigra by immunocytochemical analyses. We further identified Bach2 as a more robust inhibitor of Nrf2 responses. The effects of both Bach1 and Bach2 were dependent on their DNA-binding domains, but the DNA-binding domains did not entirely explain the differences in their relative repressor activities. Using IMR-32 neuroblastoma cells, we found differentiation into a dopaminergic neuronal phenotype resulted in increased Bach2 expression, decreased full-length Bach1 expression, increased truncated Bach1, and blunted Nrf2-mediated responses. Bach inhibitors cobalt protoporphyrin or cadmium chloride, which were more effective against Bach1 than Bach2, did not rescue Nrf2 responses. These results provide a novel mechanism for the lack of antioxidant responses and sensitivity of dopaminergic neurons to oxidative damage, in addition to proposing Bach2 as a novel drug target for the treatment of PD.

**Significance:** Oxidative stress and damage are unifying contributors to explain the progressive loss of dopaminergic neurons in Parkinson’s disease (PD). Although increased transcription of antioxidant enzymes by nuclear factor erythroid 2-related factor 2 (Nrf2) shows potential in combating oxidative damage, Nrf2 protection is mostly conferred by surrounding glia with a relative inability of neurons to mount a protective response. This work characterizes transcriptional repressors Bach1 and Bach2 that inhibit Nrf2 responses and prevent antioxidant enzyme production. This study supports Bach2 as a major factor in preventing antioxidant production in neurons and as a novel drug target to protect against PD.

## Introduction

Parkinson’s disease (PD) is the second most prevalent neurodegenerative disease and the most prevalent movement disorder. PD is characterized by the progressive loss of dopaminergic neurons in the substantia nigra pars compacta, leading to symptoms such as tremors, postural instability, and akinesia. Although several genetic mutations have been identified to be causative or increase the risk of PD, the cause or risk in idiopathic PD, ∼90% of cases, are unknown.

Oxidative stress has been identified as a central factor in both genetic and sporadic forms of PD (Dias, Junn, & Mouradian, 2013; Hwang, 2013). The most powerful defense against oxidative damage is the production of antioxidants through upregulation of transcriptional activity, designed to combat damaging oxidants such as free radicals and peroxides. The transcription factor nuclear factor erythroid 2–related factor 2 (Nrf2) provides this essential role by increasing cellular transcription of multiple antioxidant enzymes through activation of antioxidant response elements (ARE). Activation of Nrf2 results in cellular protection by increasing expression of antioxidant enzymes such as NADPH:quinone oxidoreductase 1 (NQO1), heme oxygenase 1 (HO1), and multiple components of the glutathione pathway, such as the modulatory and catalytic subunits of γ-glutamylcysteine ligase (GCLM and GCLC, respectively). The resulting increases in enzymes produce powerful antioxidants such as glutathione and reduce deleterious oxidants. Additional proteins under the transcriptional control of Nrf2 have been identified within the categories of cellular redox systems, heme metabolism, oxidoreductase activity, apoptosis and cell death, and glutathione metabolism (Chorley et al., 2012; Lacher & Slattery, 2016). Promotion of this production of antioxidants is therefore a promising mechanism to protect against the neurodegeneration of PD (Calkins et al., 2009; J. A. Johnson et al., 2008; Li, Calkins, Johnson, & Johnson, 2007).

Numerous studies have found Nrf2 activators as beneficial in models of disease, including PD, Huntington’s disease, and multiple sclerosis (D. A. Johnson & Johnson, 2015). The Nrf2 activator dimethyl fumarate (DMF) has additionally been approved by the FDA for multiple sclerosis (Wingerchuk & Carter, 2014; Xu et al., 2015). While Nrf2 activation shows some protection against neurodegeneration in animal models of PD (P. C. Chen et al., 2009; Kaidery et al., 2013; Williamson, Johnson, & Johnson, 2012; Yang et al., 2009), this response is limited in neurons with protection mostly conferred from surrounding glia (P. C. Chen et al., 2009; Lee, Calkins, Chan, Kan, & Johnson, 2003). Nrf2 is also localized to the nucleus of surviving dopaminergic neurons in patients with Parkinson’s disease (Ramsey et al., 2007), suggesting unsuccessful attempts of Nrf2 to protect these neurons against degeneration.

Comparatively to Nrf2, ARE transcriptional repressors and their roles in disease have been minimally investigated. Bach1 and Bach2 (BTB and CNC homology, basic leucine zipper transcription factor 1 and 2) are ARE-binding proteins that oppose Nrf2-mediated transcriptional activation (Oyake et al., 1996). Similar to Nrf2, Bach1 and Bach2 are members of the cap ‘n collar (CNC)-basic leucine zipper (bZIP) protein family. Bach1 and Bach2 share 38% homology, additionally containing the domains of Broad complex–Tramtrack–Bric-a-brac domain (BTB/POZ domain) at the N-terminus, and a cytoplasmic localization sequence (CLS) at the C-terminus (Hoshino et al., 2000; Kanezaki et al., 2001; Muto et al., 1998; Oyake et al., 1996). The BTB domain allows for dimer or oligomerization, which can support nuclear localization of Bach proteins (Kanezaki et al., 2001), a mechanism opposed by the CLS domain. While oxidative signals increase Nrf2 localization to the nucleus, these same stress signals can inhibit the CLS domain, promoting nuclear localization of Bach1 and Bach2, and silencing ARE-dependent transcription (Hoshino & Igarashi, 2002). Therefore, under conditions of oxidative stress, the presence of Bach1 and/or Bach2 would antagonize Nrf2 responses, thereby preventing optimal cellular protection by antioxidants.

While Bach1 is ubiquitously expressed (Blouin et al., 1998; Kanezaki et al., 2001; Ohira et al., 1998), Bach2 expression is limited to lymphocytes and neurons (Hoshino & Igarashi, 2002; Muto et al., 1998; Muto et al., 2004; Oyake et al., 1996; Roychoudhuri et al., 2013; Waclaw et al., 2017). Bach1 inhibition of Nrf2 transcriptional activity has been implicated in cardiovascular disease, pulmonary disease, and spinal cord injury (Chapple et al., 2016; Goven, Boutten, Lecon-Malas, Boczkowski, & Bonay, 2009; Goven et al., 2008; Kanno et al., 2009; Omura et al., 2005; Watari et al., 2008; Yano et al., 2006).

Additionally, Bach1 inhibitors are currently under investigation in animal models of Parkinson’s disease (Ahuja et al., 2016; Gazaryan & Thomas, 2016). However, investigation of the function of Bach2 has been mostly limited to immune disease (Noujima-Harada et al., 2017; Pazderska et al., 2016). Although Bach2 has been identified as competing with Nrf2 for the ARE transcriptional elements (Igarashi et al., 1998), only a few studies have investigated Bach2’s role in this regard (Z. Chen et al., 2013; Uittenboogaard et al., 2013).

The current work aims to characterize Bach1 and Bach2 and their potential functions in dopaminergic neurons in mediating Nrf2 antioxidant responses. We hypothesized that Bach1 and/or Bach2 would inhibit the activity of Nrf2, resulting in a lack of protective antioxidant enzyme expression. We found expression of Bach1 and Bach2 in the substantia nigra and in model cell systems determined the relative repressor abilities of Bach1 and Bach2. We further correlated changes in Bach1 and Bach2 expression with the effects of neuronal differentiation on Nrf2-mediated responses. The current results highlight the potent inhibitory abilities of Bach2 in opposing Nrf2-mediated antioxidant enzyme production. This work supports that Bach2 responses may prevent dopaminergic neurons from adequately producing antioxidants that could confer a protective response against PD and identify Bach2 as a novel drug target for PD.

## Materials and Methods

### Cell culture

HEK293 cells (ATCC Cat# CRL-1573, RRID:CVCL_0045) were maintained in Dulbecco’s Modified Eagle’s Medium (DMEM; Invitrogen) supplemented with 10% fetal bovine serum (FBS) and 100 U/ml penicillin/ 100 µg/ml streptomycin. IMR-32 neuroblastoma cells (ATCC Cat# CCL-127) were maintained in DMEM/Ham’s F12 (1:1) with 15mM HEPES (Invitrogen) supplemented with 10% FBS and 100 U/ml penicillin/ 100 µg/ml streptomycin. IMR-32 neuroblastoma were differentiated for 15-16 days using the above media supplemented with 5-Bromo-2’-deoxyuridine (BrdU; 2.5 µM; Sigma) and N6, 2’-O-dibutyryladenosine 3’,5’-cyclic monophosphate sodium salt (db-cAMP; 1 mM; Sigma). Media was replaced every 2-3 days for a minimum of 15 days. LUHMES neuroblastoma cells (ATCC Cat# CRL-2927) were maintained DMEM/F12 (ATCC) supplemented with N-2 supplement (1%; Invitrogen) and human recombinant basic fibroblast growth factor (40 ng/ml; R&D Systems) on plates pre-coated with poly-l-ornithine (50 µg/ml; Sigma) and fibronectin (1 µg/ml; Sigma). LUHMES neuroblastoma were differentiated using media supplements with tetracycline (1 µg/ml; Sigma), db-cAMP (1 mM), and glial derived neurotrophic factor (2 ng/ml; R&D Systems). One day after the initiation of differentiation, cells were split and re-seeded onto fresh coated plates. Media was replaced every 2-3 days for a minimum of 6 days. All cells were maintained in a trigas incubator containing 5% CO_2_ and 5% O_2_, supplemented with N_2_, unless otherwise specified. Indicated drug treatments were performed for approximately 24h prior to the harvesting of cells. Mono-methyl fumarate (MMF), cobalt protoporphyrin (CoPP), and cadmium chloride (CdCl_2_) were each obtained from Sigma. Drug treatments were matched by dimethyl sulfoxide (DMSO) in the control condition. Higher concentrations of DMSO did not produce a similar response to no-treatment control conditions, so the DMSO content per well was limited to 0.05%. Experiments with higher concentrations of DMSO were omitted from analysis.

Plasmids and cell transfections: Mammalian expression vectors containing cDNA for full-length human Bach1 (in pCMV6-XL5; SC308292), Bach2 (in pCMV6-XL4; SC112552), Nrf2 (in pCMV6-XL5; SC116283), and empty vector (pCMV6-Entry; pCMV) were obtained from Origene.

For transfection of HEK293 cells, plates were pre-coated with poly-*D*-lysine (10 µg/ml; Sigma). Cells were plated to be at approximately 25% confluency during the day of transfection. Transfections were performed by calcium phosphate precipitation, as previously described (Waxman & Giasson, 2010). Transfection of IMR-32 neuroblastoma cells was performed, as above with the following modifications: media was replaced with OptiMem (Invitrogen) prior to transfection, and 60 µl (rather than 75 µl) of final transfection solution was added to each 35mm well.

Approximately 16 hours after transfection, the calcium phosphate precipitant was washed off the cells two-times with PBS, and then the media was replaced (to DMEM or DMEM/F12, as above). HEK293 cells were harvested 48-72 hours after transfection.

### Creation of Bach1 and Bach2 deletion and hybrid cDNA

Primers were created to PCR the indicated regions of Bach1 or Bach2, adding an NheI digestion site at the 5’ and an XhoI site at the 3’ end of the PCR. In some cases the natural XhoI site contained in the vector construct was used. PCR products were ligated into the pcDNA3.1 vector (Invitrogen). For Bach1 delta-BTB, the following forward and reverse primers were used to place a methionine and start the construct with D132: F - tctgaaatgctagcttatggactccactgcagaccagcaagaat; R - ctgccaaaactcgaggagaaaattagatggtttgaaggaa. For the Bach2 delta-BTB, the following forward and reverse primers were used to place a methionine and start the construct with N135: F - ttcctgcaga<u>GcTagc</u>tc Atgaacagtgaggatggcctgttt; R - tgt gaa ggt aac tat cac tcc tgc tcg aga aga gga. For Bach1 delta-CLS, the following forward and reverse primers were used to place a stop codon after D723: F – TGATAATTAGAA<u>GCtaGC</u>TTTCCACTGAACTTCCCGACAACA; R – tat cag tcat <u>ctc gag</u> tca tta atc tgag atccc acc act. For Bach2 delta-CLS, the following forward and reverse primers were used to place a stop codon after T819: F - atcctaagactcgaag<u>gcTagc</u>acaggacctggaaa aattac; R- ctggcagaa ctc gag cta ggtcactgtttggctcctgggttcaaga.

For deletions or modifications to the DNA binding domains, PCRs were performed by overlapping extension reactions, such that two initial PCRs were performed with 1) a general forward and a reverse containing half of one side of the deleted sequence, and 2) with a general reverse and a forward containing the other side of the sequence, but with the reaction 1’s reverse being the reverse complement to reaction 2’s forward, above. After the performance of the two initial PCR reactions, the two PCR products were purified and then PCRed together using the general forward and reverse primers, ligating the two products together. For the Bach1 delta-CNC-bZIP construct, the following overlapping extension primer sequence was used to delete amino acids 527-623: ACAAGAATGTGAGGTAAAACTG-GTTTGTAAAGAAGCAGCTCT. For the Bach2 delta-CNC-bZIP construct, the following overlapping extension primer sequence was used to delete amino acids 616-712: aggggccaggaggtaaaactt-gtttgccgagacatccaga. The same overlapping extension PCR technique was used to create hybrid cDNAs; however, an additional initial PCR was performed, and three products were used in the final PCR reaction. To create the Bach1 cDNA containing the Bach2 CNC-bZIP, the following primer was used to extend from amino acid 526 of Bach1 to amino acid 616 of Bach2: ACAAGAATGTGAGGTAAAACTG-ccttttcctgtagatcaaat. The following primer was used to extend from amino acid 712 of Bach2 back to amino acid 624 of Bach1: tctcctgcctttcccaggaa-GTTTGTAAAGAAGCAGCTCT. To create the Bach2 cDNA containing the Bach1 CNC-bZIP, the following primer was used to extend from amino acid 615 of Bach2 to amino acid 527 of Bach1: aggggccaggaggtaaaactt-CCATTCAATGCACAACGGATA. The following primer was used to extend from amino acid 623 of Bach1 back to amino acid 713 of Bach2: TAACTGGACTTTGCCAGAAA-gtttgccgagacatccaga. For overlapping extension reactions, only the forward primers are provided.

To create the Bach-BTB only myc-tagged cDNAs, the BTB domain regions of each were PCRed from the original vectors from Origene, and then ligated into the pCMV6-Entry vector using HindIII and XhoI restriction enzymes. The following primers were used to create Bach1-BTB: forward: ATGATAATTAGAAGCTTGCTTTCCACTGAACT, reverse: agc att ttt ttc ttg ggc act cga gct ggt ctg cag tgg agt. The following primers were used to create Bach2-BTB: forward: agttccctgcatcctaagcttcgaaggcagcaca, reverse: AGCATCCTTCCGGCACTCGAGCAGGCCATCCTCACT.

All primer sequences listed are in the 5-prime to 3-prime direction. All sequences once cloned into the final vector were confirmed by Sanger sequencing through the entire open reading frame.

### Western blot analysis

Samples were solubilized in 1.5X Laemmli buffer (75 mM Tris-HCl, pH 6.8, 3% SDS, 15% glycerol, 3.75 mM EDTA, pH 7.4), followed by heating to 100°C for 10 min. Protein concentrations were determined using Bicinchoninic Acid Assay (ThermoFisher/ Pierce), and then samples were incubated with dithiothreitol and bromphenol blue and heated to 100°C for 10 min prior to Western blot analysis. Protein samples were resolved by SDS-PAGE on 7% or 12% gels, depending on molecular weight for the proteins of interest, followed by electrophoretic transfer onto nitrocellulose membranes. Membranes were blocked in Tris buffered saline (TBS) with 5% dry milk, and incubated overnight with antibodies specific to NQO1 (1:500; Abcam, ab34173 or 1:250; Cell Signaling, clone A180), HO1 (1:500; Millipore, 374090), GCLM (1:5000; Millipore, MABS32), GCLC (1:1000; Abcam, ab190685 or 1:300; Millipore clone 3H1), glutathione-*S-*reductase (GSR; 1:1000; Abcam, ab124995), Bach1 (1:500; Bethyl Labs, 057A), Bach2 (1:2000; Sigma-Aldrich Cat# SAB2100201, or Atlas Antibodies Cat# HPA051384, purchased from Sigma-Aldrich), Nrf2 (1:250; Santa Cruz, H-300), tyrosine hydroxylase (TH; 1:5000; Millipore, AB152), β-actin (actin; 1:1000 Sigma, A2103), and glyceraldehyde 3-phosphate dehydrogenase (GAPDH; 1:1000; Cell Signaling, 3683) diluted in TBS / 5% dry milk or TBS/ 5% bovine serum albumin (BSA). See Table 1 for additional antibody information. Each incubation was followed by goat anti-mouse conjugated horseradish peroxidase (HRP) (Cell Signaling), goat anti-rabbit conjugated HRP (Cell Signaling), or donkey anti-goat conjugated HRP (Santa Cruz), and immunoreactivity was detected using chemiluminscent reagent (West Pico, ThermoFisher-Pierce or WesternBright ECL, Advansta, Menlo Park, CA) and image capture with an ImageQuant LAS4000 system and software (GE).

**Table 1.**
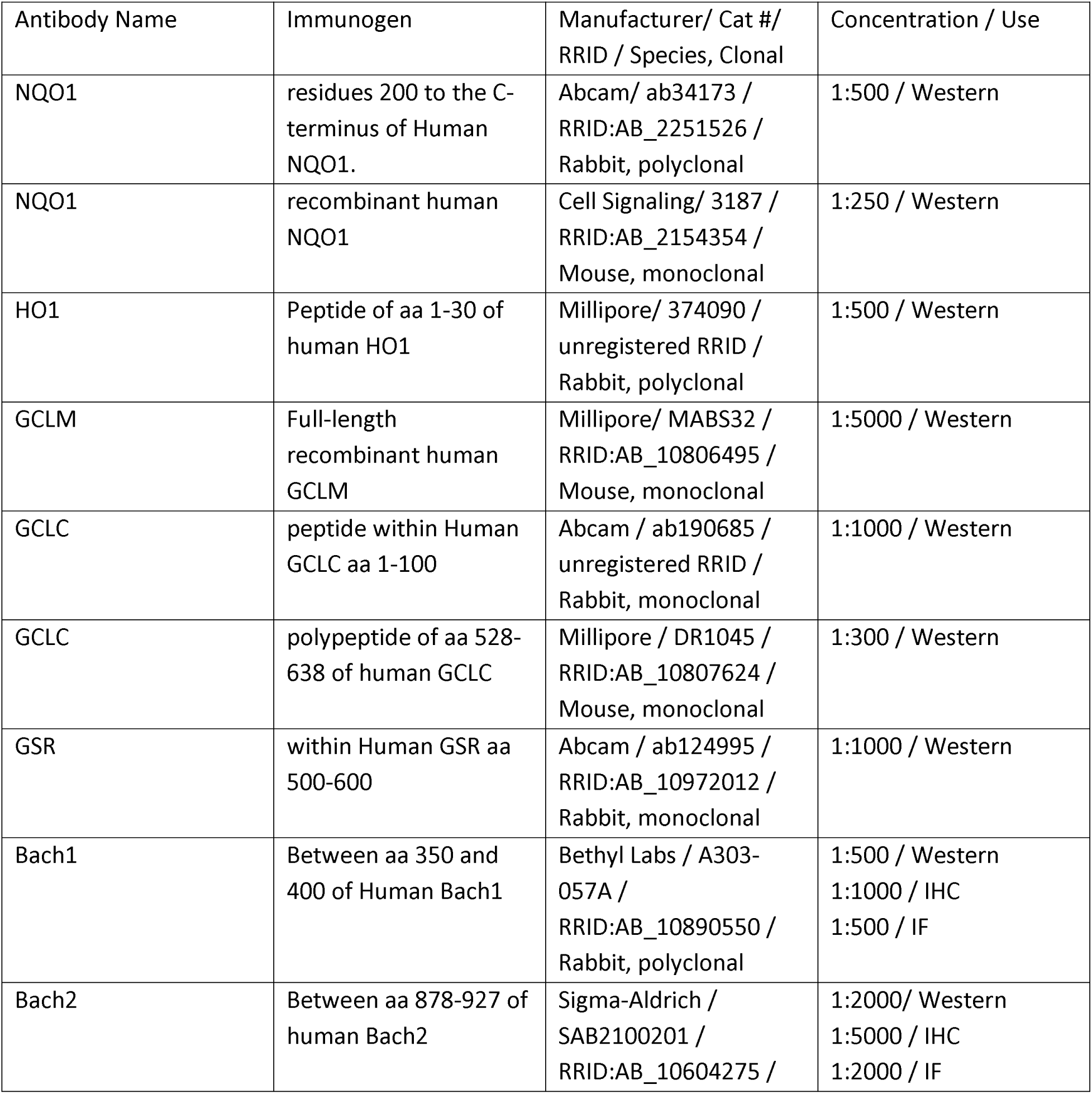
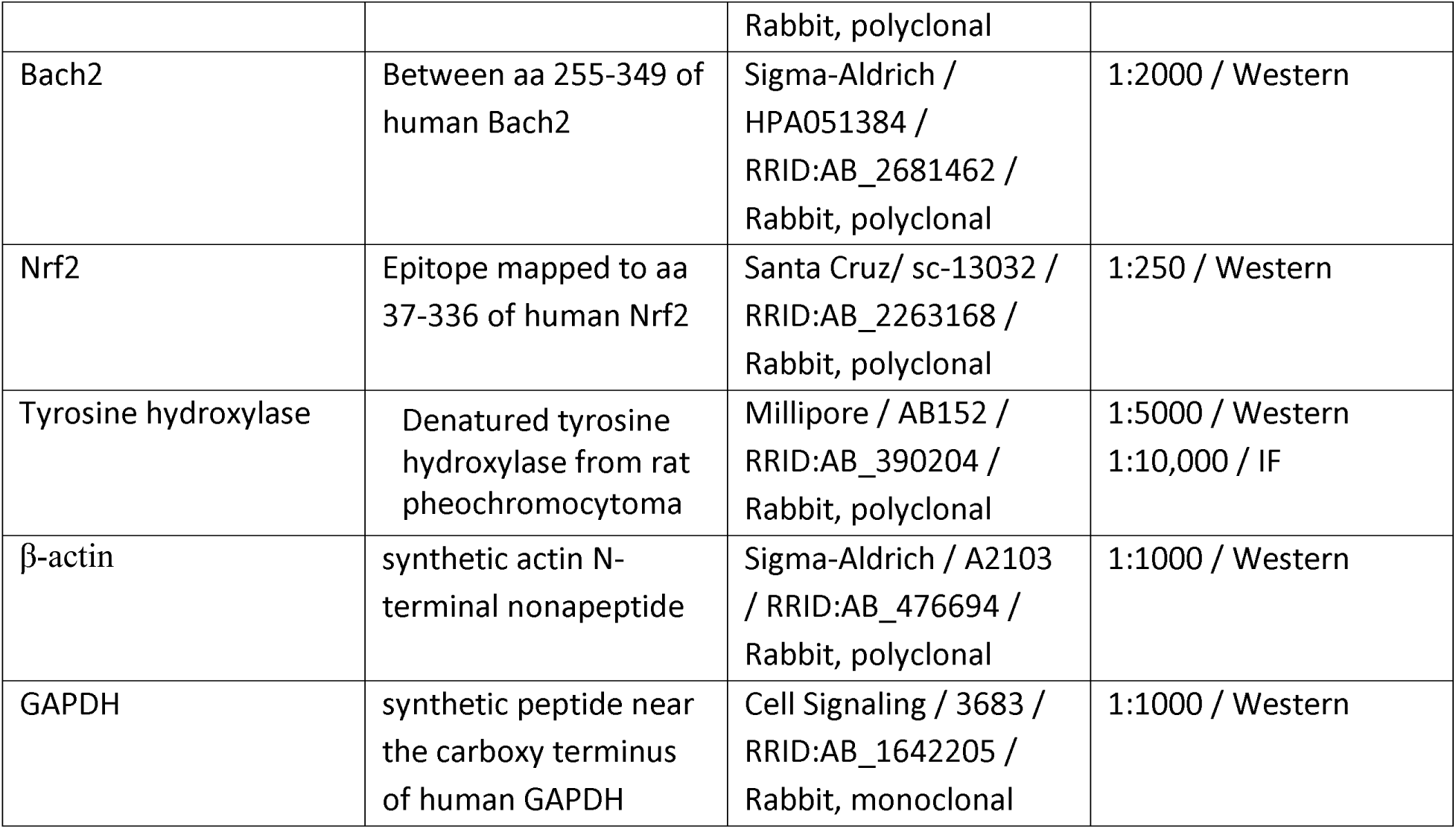
Antibody information for each antibody used during the course of this study. All antibodies used were commercially available, as indicated by the manufacturer and catalog number (Cat #) indicated. Immunogen information was provided as available. In some cases the exact residues were not made available by the manufacturer. Abbreviations: aa = amino acid residues; IHC = immunohistochemistry; IF = immunofluorescence.

| Antibody Name | Immunogen | Manufacturer/ Cat #/<br>RRID / Species, Clonal | Concentration / Use |
| --- | --- | --- | --- |
| NQO1 | residues 200 to the C-terminus of Human NQO1. | Abcam/ ab34173 /<br>RRID:AB_2251526 /<br>Rabbit, polyclonal | 1:500 / Western |
| NQO1 | recombinant human NQO1 | Cell Signaling/ 3187 /<br>RRID:AB_2154354 /<br>Mouse, monoclonal | 1:250 / Western |
| HO1 | Peptide of aa 1-30 of human HO1 | Millipore/ 374090 /<br>unregistered RRID /<br>Rabbit, polyclonal | 1:500 / Western |
| GCLM | Full-length recombinant human GCLM | Millipore/ MABS32 /<br>RRID:AB_10806495 /<br>Mouse, monoclonal | 1:5000 / Western |
| GCLC | peptide within Human GCLC aa 1-100 | Abcam / ab190685 /<br>unregistered RRID /<br>Rabbit, monoclonal | 1:1000 / Western |
| GCLC | polypeptide of aa 528-638 of human GCLC | Millipore / DR1045 /<br>RRID:AB_10807624 /<br>Mouse, monoclonal | 1:300 / Western |
| GSR | within Human GSR aa 500-600 | Abcam / ab124995 /<br>RRID:AB_10972012 /<br>Rabbit, monoclonal | 1:1000 / Western |
| Bach1 | Between aa 350 and 400 of Human Bach1 | Bethyl Labs / A303-057A /<br>RRID:AB_10890550 /<br>Rabbit, polyclonal | 1:500 / Western<br>1:1000 / IHC<br>1:500 / IF |
| Bach2 | Between aa 878-927 of human Bach2 | Sigma-Aldrich /<br>SAB2100201 /<br>RRID:AB_10604275 / | 1:2000/ Western<br>1:5000 / IHC<br>1:2000 / IF |
|  |  | Rabbit, polyclonal |  |
| Bach2 | Between aa 255-349 of human Bach2 | Sigma-Aldrich / HPA051384 / RRID:AB_2681462 / Rabbit, polyclonal | 1:2000 / Western |
| Nrf2 | Epitope mapped to aa 37-336 of human Nrf2 | Santa Cruz/ sc-13032 / RRID:AB_2263168 / Rabbit, polyclonal | 1:250 / Western |
| Tyrosine hydroxylase | Denatured tyrosine hydroxylase from rat pheochromocytoma | Millipore / AB152 / RRID:AB_390204 / Rabbit, polyclonal | 1:5000 / Western<br>1:10,000 / IF |
| β-actin | synthetic actin N-terminal nonapeptide | Sigma-Aldrich / A2103 / RRID:AB_476694 / Rabbit, polyclonal | 1:1000 / Western |
| GAPDH | synthetic peptide near the carboxy terminus of human GAPDH | Cell Signaling / 3683 / RRID:AB_1642205 / Rabbit, monoclonal | 1:1000 / Western |

### Antibody Characterization

All antibodies were validated by Western blot analyses. Antioxidant enzyme antibodies were verified by recognition of correct molecular weight and increases in expression with Nrf2 activation. GCLM and GCLC were additionally characterized through overexpression in HEK293 cells. Bach1 and Bach2 antibodies were verified with overexpression in HEK293 cells and shRNA (plasmids purchased from Origene) knockdown of endogenous protein in HEK293 cells for Bach1 and differentiated IMR-32 cells for Bach2 (data not shown). Nrf2 antibodies were validated by overexpression of Nrf2 in HEK293 cells and increase in Nrf2 expression with use of Nrf2 chemical activators in HEK293 cells and IMR-32 cells.

NQO1 and GCLC antibodies were switched during the course of this study to Abcam antibodies when new lots of original antibodies did not produce the same specificity as the older lots (Cell Signaling and Millipore, respectively). NQO1 (Cell Signaling) was used for Figures 4 & 7, and NQO1 (Abcam) was used for the remainder of the study. GCLC (Millipore) was used for Figures 4 & 5, and GCLC (Abcam) was used for the remainder of the study. The newer antibodies were compared against previous immunoreactivities of the original antibodies in this study under the same HEK293 transfection conditions prior to switching for the remainder of the study. Both Bach2 antibodies provided similar immunoreactivities in samples. For representative immunoblots, Bach2 HPA051384 was used for Figures 5 & 11, and Bach2 SAB2100201 was used for all other Figures and quantitative analysis.

### Coimmunoprecipitation

Coimmunoprecipitation was performed as previously described (Waxman, Baconguis, Lynch, & Robinson, 2007) with the following substitutions: Protein A/G agarose (Santa Cruz Biotechnology, Santa Cruz, CA) was used to bind 3ug of antibody. Normal rabbit IgG (Santa Cruz) was used as the comparator antibody for IP. Transfections for IP were performed with combinations of Bach1, Bach2, Bach1-BTB (B1-BTB), Bach2-BTB (B2-BTB), or myc control (pCMV6-entry plasmid). Final precipitation from beads were completed by heating to 100°C in 1.5X Laemmli buffer for 10min.

### Immunohistochemistry

Formalin-fixed paraffin-embedded human substantia nigra samples were provided by Harvard Brain Tissue Resource Center (HBTRC; Cambridge, MA). This research was covered under the HBTRC ethical protocols and Partners Human Research Committee approvals. Samples were de-identified by HBTRC prior to being provided to the present author/ institution. Samples were provided based on limited availability of paraffin blocks, and matched between PD and control samples as well as possible by age and gender. Both genders were included in a random fashion, based on availability of 7 Parkinson’s disease patients, and 7 control patients (Table 2). A diagnosis was provided by Harvard Brain Tissue Resource Center, and pathology reports were reviewed by E. Waxman for consistency with diagnosis. Parkinson’s disease patients with notes of dementia, tau pathology, and/or Abeta pathology were indicated as Parkinson’s disease with dementia (PDD). All samples were deemed diagnostically appropriate for analysis. Samples were sectioned at 10 µm thickness on to charged slides. Paraffin was dissolved in xylene, and sections were serially re-hydrated in decreasing concentrations of ethanol. After a 10 min water rinse, sections underwent heat-induced epitope retrieval with 1X citrate buffer, pH 6.0 (Sigma) for a 7 min pressure cooker setting, followed by 20 min depressurization. Samples were blocked for 1 h in Tris, pH7.5 containing rabbit serum, and then incubated overnight with Bach1 (1:1000; Bethyl Labs, 057A) or Bach2 (1:5000; Sigma-Aldrich Cat# SAB2100201) antibody diluted in blocking solution, or blocking solution alone, for no primary condition. Approximately 16 h later, samples were washed, then incubated with goat anti-rabbit biotinylated secondary antibody, followed by the ABC-method (Vector Labs) conjugated to alkaline phosphatase. The enzyme was developed using Vector Red substrate and samples were counterstained with hematoxylin. Sections were then dehydrated through increasing concentrations of ethanol, cleared with xylene, and mounted with cytoseal (ThermoFisher).

**Table 2.** Demographics of patient samples used for anti-Bach1 and anti-Bach2 immunoreactivity analysis in the substantia nigra. Abbreviations: PMI = post-mortem interval; PDD = Parkinson’s disease with dementia; M = male; F = female.

| Case # | Diagnosis | Age | Sex | PMI |
| --- | --- | --- | --- | --- |
| 1 | PDD | 79 | F | 21.12 |
| 2 | PDD | 82 | M | 18.08 |
| 3 | PDD | 84 | F | 18.92 |
| 4 | Parkinson | 79 | F | 4.95 |
| 5 | Parkinson | 73 | F | 24.00 |
| 6 | Parkinson | 76 | M | 12.95 |
| 7 | Parkinson | 81 | F | 13.8 |
| 8 | Control | 55 | F | 21.66 |
| 9 | Control | 67 | M | 23.12 |
| 10 | Control | 58 | F | 19.38 |
| 11 | Control | 65 | M | 20.92 |
| 12 | Control | 73 | M | 18.02 |
| 13 | Control | 85 | M | 20.83 |
| 14 | Control | 56 | M | 17.32 |

### Tissue immunofluorescence

Immunofluorescence on tissue was performed after re-hydration and heat-induced epitope retrieval, as above. Samples were blocked with Tris, pH 7.5 containing 5% dry milk, and antibodies were incubated overnight at 4°C in a humidifying chamber. Antibodies Bach1 (1:500) or Bach2 (1:2000) were co-incubated with TH (1:10,000, Sigma, T1299). After primary incubation and washes in Tris, pH7.5, fluorescent secondary antibody incubation with goat anti-mouse Alexa 594 and goat anti-rabbit Alexa 488 (each at 1:500; Invitrogen) was performed for 2 h at room temperature. For all sample types, nuclei were counterstained with Hoechst trihydrochloride trihydrate 3342 (Invitrogen), and coverslips were mounted using Fluoromount-G (Southern Biotech, Birmingham, AL).

### Microscopy

Light microscopy was performed using an inverted light/ fluorescent microscope with a 10X and 20X objective (Amscope) and a Sony NEX-3 camera mounted for image capture. Confocal microscopy was performed with a Zeiss Axiovert 200M inverted confocal microscope mounted with a Zeiss LSM520 META NLO digital camera utilizing Zeiss LSM510 META V3.2 confocal microscope software (Zeiss, Thornwood, NY). Confocal images were captured with 63x oil optics. Representative images of confocal microscopy of brain tissue were of a single Z-plane of <1.5 µm.

### Quantitative analysis

Western blot data quantified densitometry was performed by FUJI/ ImageJ software (NIH, Bethesda, MD). Expression levels were standardized to densitometry of β-actin. All data were further standardized to the control (vehicle or vector only) or the Nrf2 expression (activation) condition, except where otherwise indicated. Statistical analyses were performed using RStudio (Version 1.1.456). One-way ANOVA analyses were performed for each antibody using aov(standardized_densitometry ∼ condition) with *p*-values for the *F* statistic reported. Post-test analyses (two-way *t*-tests) were performed by pairwise.t.test(standardized_densitometry, condition, p.adjust = “none”, pool.sd=F, var.equal=T), followed by p.adjust(pairwise_score, method = “bonferroni”, n = comparisons) selected for targeted comparisons, with comparison number limited to direct comparisons within each antibody. Exceptions include those of unequal variances (omitting var.equal argument) and comparisons against the standardized conditions, which were performed by one-sample *t*-tests were performed with t.test() and mu = 100 or 0 (the value of the control/ standardized condition), followed by p.adjust() for Bonferroni corrections. For two-way or higher factor ANOVA analyses, *F* tests were performed using aov(standardized_densitometry ∼ factor1 * factor2 (etc)) for each factor and potential interaction of factors. Statistically significant factors and interactions were further interrogated as group-wise comparisons by performing t.test() as paired = True to directly compare factor differences. Subsequent post-hoc tests were performed as indicated above by pairwise.t.test() or t.test(), followed by Bonferroni corrections. Each experiment was performed a minimum of three independent times, containing paired controls for each experimental condition. Random numbers provided by Harvard Brain Resource Center was used for blinding the brain tissue samples. Evaluation of double-immunofluorescence was performed by capture of at least 3 random TH-positive fields, followed by blinded semi-quantitative intensity analyses using ImagePro Premier software (Media Cybernetics, Rockville, MD).

## Results

### Bach1 and Bach2 expression in the human substantia nigra

Bach1 expression is reported to be ubiquitous (Blouin et al., 1998; Kanezaki et al., 2001; Ohira et al., 1998), and Bach2 expression is limited to certain cell types such as lymphocytes and neurons (Hoshino & Igarashi, 2002; Muto et al., 1998; Oyake et al., 1996; Roychoudhuri et al., 2013; Waclaw et al., 2017). We hypothesized that both Bach1 and Bach2 would be expressed in the human substantia nigra, potentially diminishing Nrf2’s ability to protect against neurodegeneration. We performed immunohistochemical analysis on human substantia nigra tissue, using antibodies specific for human Bach1 or Bach2 and the Vector Red enzyme (**Fig 1**). Red immunostaining and 564nm fluorescence were identified in the tissue with either anti-Bach1 or anti-Bach2 antibodies, co-localizing with the melanin-pigmented (brown) substantia nigra dopaminergic neurons. Anti-Bach1 antibodies also produced an overall increase of fluorescence of the neuropil, consistent with ubiquitous expression of Bach1, with a subset of dopaminergic neurons presenting with more robust staining (arrowheads).

**Figure 1.**
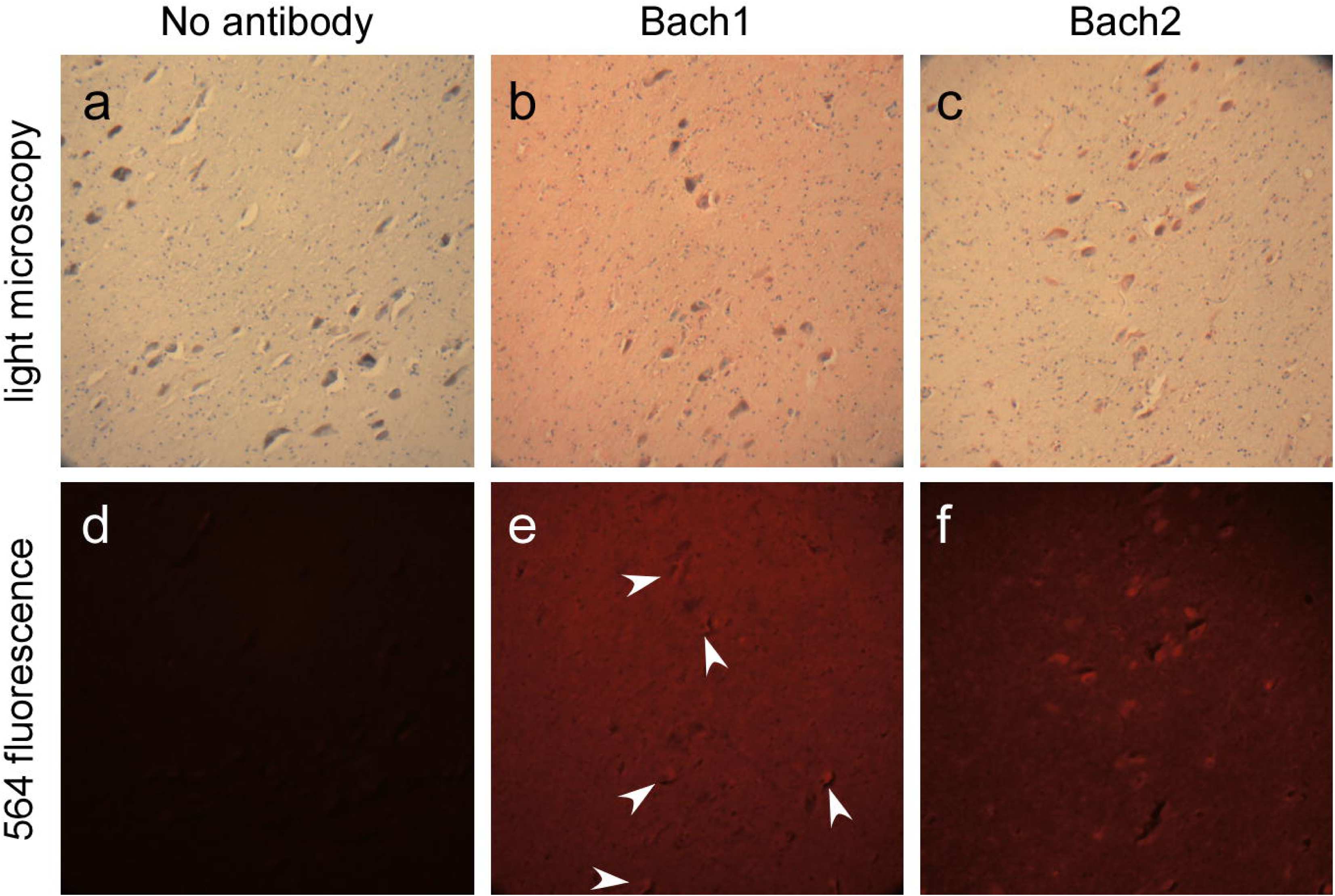
Bach1 and Bach2 expression in substantia nigra neurons of the human brain by immunohistochemistry. Immunohistochemistry was performed using the ABC method conjugated to alkaline phosphatase and Vector Red enzyme substrate on human substantia nigra samples. Panels a-c were captured with light microscopy, and panels d-f were captured of the same field, concurrently, but with 564nm fluorescence. Post-mortem, human substantia nigra brain samples stained with antibodies specific to human Bach1 (b,e) or Bach2 (c,f). Absence of primary antibody yielded only background neuromelanin staining (brown) of dopaminergic neurons in the substantia nigra (a) and a lack of 564nm fluorescence (d). (b,e) Anti-Bach1 immunoreactivity was observed throughout the tissue in white matter, processes, and neurons with some additional intensity of reactivity in neurons that co-localize with neuromelanin staining (e, arrowheads). (c,f) Anti-Bach2 immunoreactivity was observed co-localizing with neuromelanin-positive neurons.

We further performed double-immunofluorescence and confocal microscopy analysis between antibodies specific to Bach1 or Bach2 and tyrosine hydroxylase (TH), the rate-limiting enzyme in the production of dopamine and marker for dopaminergic neurons (**Figs 2 and 3**). Both Bach1 and Bach2 were expressed in TH-positive neurons of both PD patients and controls (demographics of samples analyzed are indicated in Table 2). Using ImagePro Premier software to analyze expression in TH-positive neurons, Bach1 expression was 100%. This analysis may have been partially confounded by the Bach1 staining in the neuropil (glia/ processes), despite the limited Z-plane in the confocal analysis (<1.5 µm).

**Figure 2.**
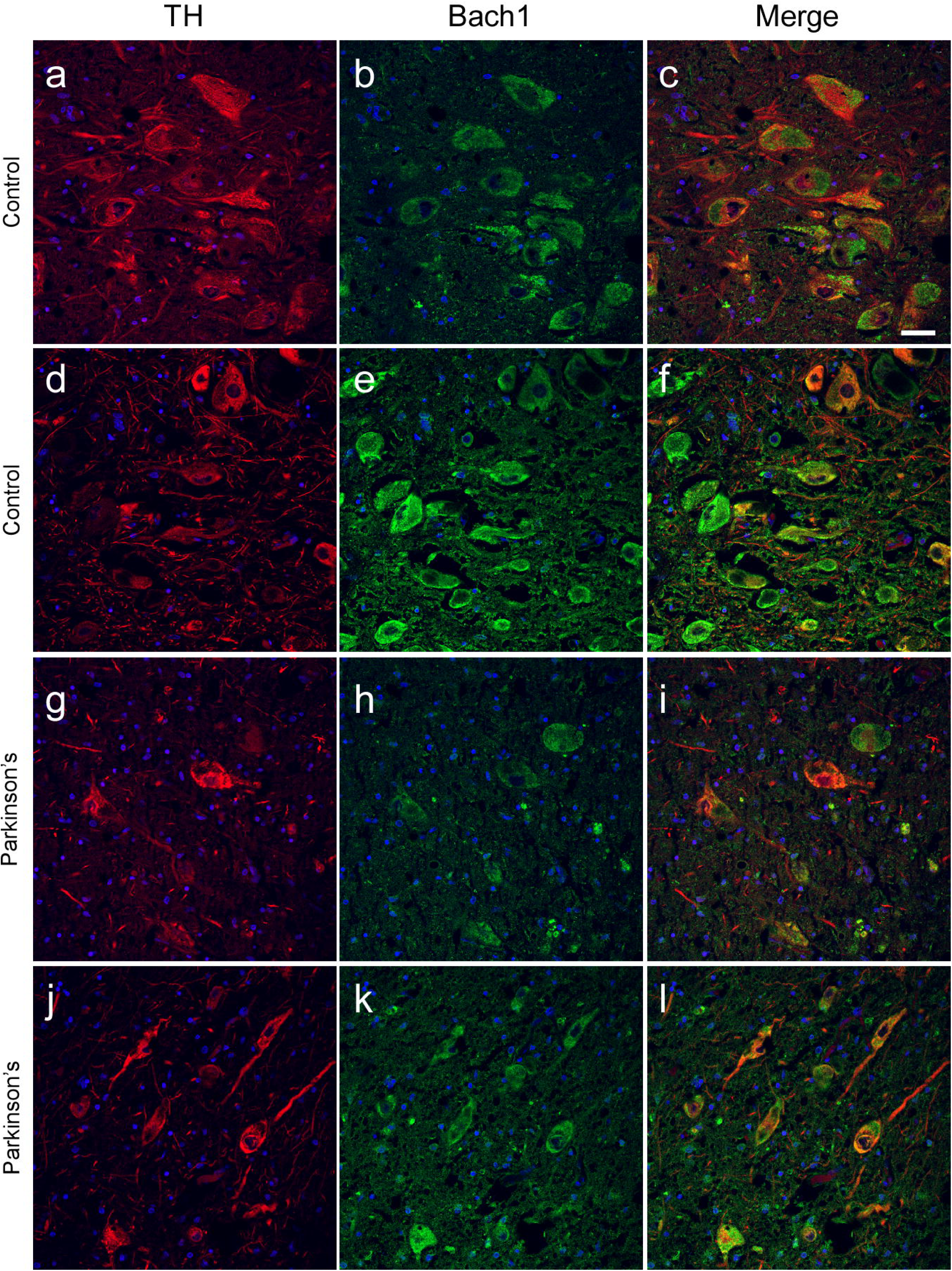
Double-immunofluorescence of Bach1 and tyrosine hydroxylase (TH) in the human substantia nigra. Representative confocal microscopy images from the human substantia nigra of Control and Parkinson’s disease (PD) samples, immunostained with antibodies specific to TH (a,d,g,j; red) and Bach1 (b,e,h,k; green). Samples were counterstained with Hoechst for nuclear visualization. Evaluation of Bach1 immunoreactivity identified generalized staining throughout the tissue with more robust staining in some neuronal cell bodies. Intensity levels of Bach1 immunoreactivity were not significantly different between control and PD samples. Scale bar = 40μm.

**Figure 3.**
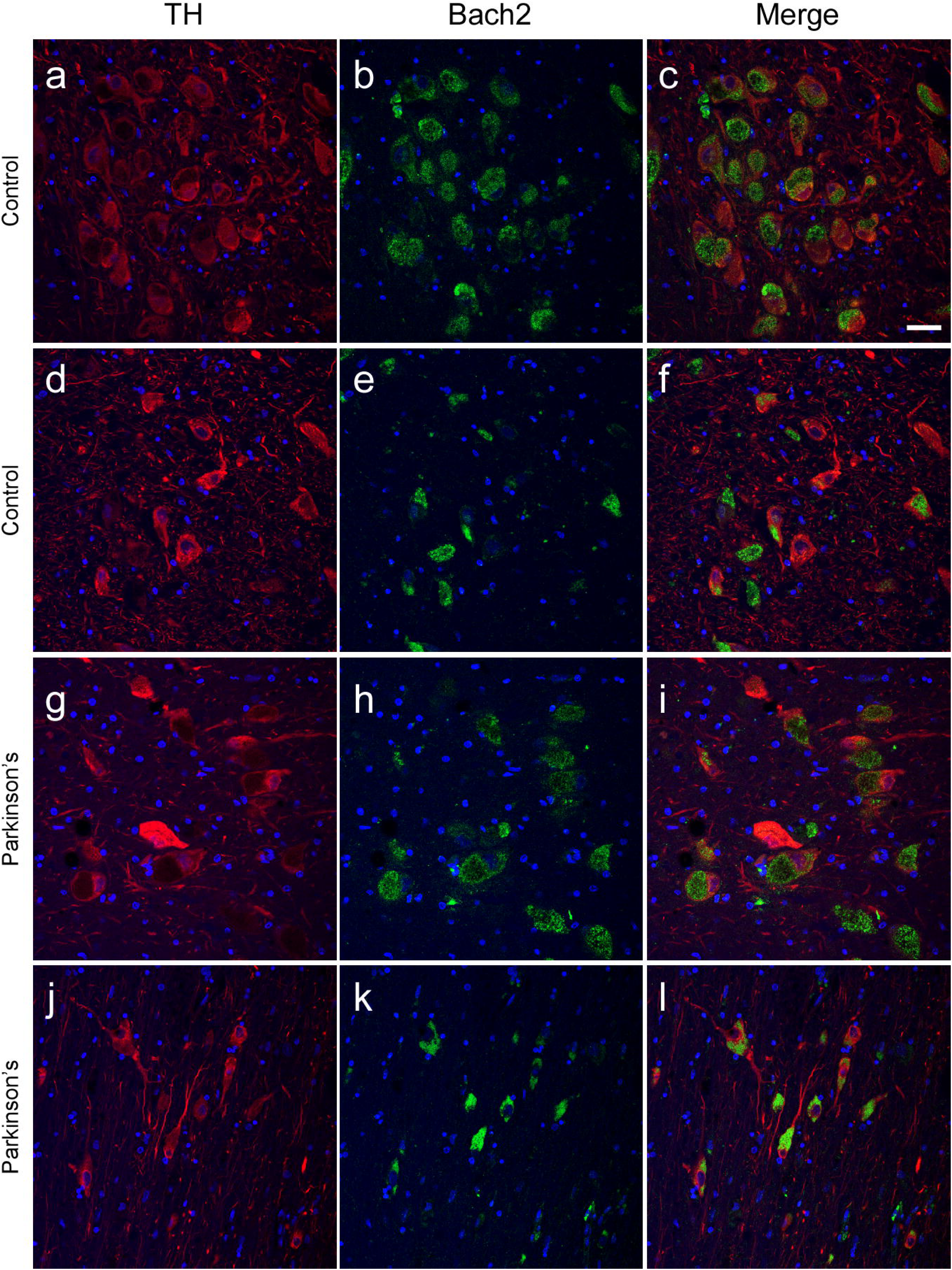
Double-immunofluorescence of Bach2 and tyrosine hydroxylase (TH) in the human substantia nigra. Representative confocal microscopy images from the human substantia nigra of Control and Parkinson’s disease (PD) samples, immunostained with antibodies specific to TH (a,d,g,j; red) and Bach2 (b,e,h,k; green). Samples were counterstained with Hoechst for nuclear visualization. Bach2 was predominantly observed in the cellular soma, often peri-nuclear localization, in the cells as those TH+, but not overlapping in sub-cellular localization (c,f,i,l). Rare non-neuronal cells or processes were observed with anti-Bach2 immunoreactivity. Evaluation of 14 samples (7 control and 7 Parkinson’s disease; PD) show 95% of TH+ neurons also contain anti-Bach2 immunoreactivity. Intensity levels of anti-Bach2 immunoreactivity in TH+ neurons and localization of anti-Bach2 immunoreactivity were not significantly different between control and PD samples. Evaluations were made from 4-7 random fields per sample. Scale bar = 40μm.

Bach2 expression was mostly limited to the cellular soma of TH-positive neurons, but rarely showing sub-cellular co-localization with TH itself. Bach2 expression was identified in 91.9±4.6% of TH-positive neurons in PD samples and 93.4±3.8% of control samples. No significant differences were identified in the PD patients versus controls. The only difference observed between the neurons was the stereotypical loss of dopaminergic neurons in the substantia nigra of PD patients, and therefore overall fewer neurons to analyze in PD patient samples.

### Relative inhibitory properties of Bach1 and Bach2

We then examined relative inhibitory potentials of Bach1 and Bach2 by performing a comparative analysis of Bach1 and Bach2 effects on Nrf2 transcriptional activity using model cell systems. We overexpressed Bach1 or Bach2 in conjunction with Nrf2 in HEK293 cells and performed Western blot analysis using antibodies specific for Nrf2-controlled antioxidant enzymes, NQO1, HO1, GCLM, and GCLC (**Fig 4**). Statistically significant difference in antioxidant enzyme expression levels were identified between the transfection conditions, with the exception of GCLC (Table 3). Co-expression of Bach1 only significant inhibited Nrf2-promoted HO1 expression levels. However, co-expression of Bach2 with Nrf2 reduced all antioxidant enzyme expression levels back to basal levels.

**Figure 4.**
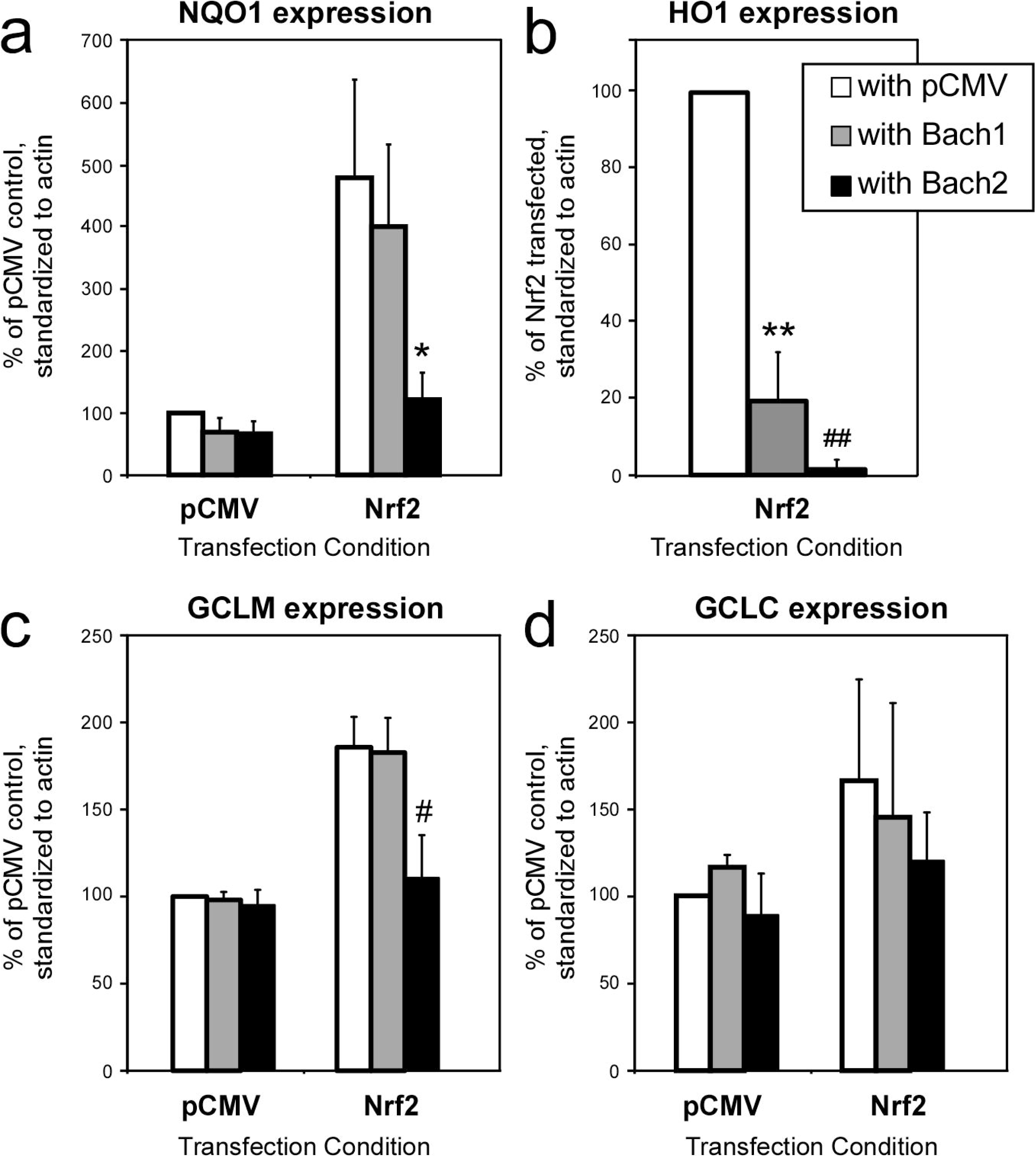
Overexpression of Nrf2 and Bach proteins in HEK293 cells each effect antioxidant enzyme expression. Quantitative densitometry of Western blot data from HEK293 cells transiently co-transfected with combinations of pCMV (empty vector control) or Nrf2 cDNA with pCMV, Bach1 cDNA, or Bach2 cDNA. Each cDNA represented 1/2 of the DNA in the transfection mix. Cells were maintained at 5% CO_2_ and 5% O_2_ and harvested 48-56 hours after transfection. Western blot analyses were performed with antibodies specific to (a) NQO1, (b) HO1, (c) GCLM, or (d) GCLC. Antibody immunoreactivity was standardized to actin, and provided as a percent of the antioxidant to actin ratio for the pCMV control for a,c,d and provided as a percent of the ratio for the Nrf2/ pCMV condition for b. Of note, no HO1 immunoreactivity was identified in the absence of Nrf2 overexpression in HEK293 cells. Statistically significant differences were identified between Nrf2/ pCMV versus Nrf2/ Bach2 co-expression for NQO1, GCLM (*, *p*=0.04 and #*p*=0.03, respectively, by two-way t-test), and HO1 (**, *p*=0.01; ##, *p*=0.0006 by one-sample t-test, each with Bonferroni corrections). Data represent average ± SD of three independent experiments.

**Table 3.** Statistical significance of differences between Nrf2, Bach1, and Bach2 cDNA transfections in HEK293 cells. One-way ANOVA tests were performed comparing all transfection conditions, followed by post-hoc two-way student t-tests, or for HO1 two-way one-sample t-test, with Bonferroni corrections (comparisons=2). pCMV versus pCMV/Nrf2 comparisons were performed by one-sample t-tests, except HO1, which was not done (ND) due to the lack of signal in the pCMV condition. Statistically significant *p*-values are shown in bold.

| Antibody | ANOVA | *P*-values – post-hoc t-tests with Bonferroni corrections |  |  |
| --- | --- | --- | --- | --- |
|  |  | pCMV vs pCMV/Nrf2 | pCMV/ Nrf2 vs Bach1/Nrf2 | pCMV/Nrf2 vs Bach2/Nrf2 |
| NQO1 | $F_{5,12} = 13.4$<br>$p = 0.0001$ | 0.053 | 1.0 | **0.04** |
| GCLM | $F_{5,12} = 24$<br>$p < 0.0001$ | **0.01** | 1.0 | **0.03** |
| GCLC | $F_{5,12} = 1.62$<br>$p = 0.23$ | 0.57 | 1.0 | 0.57 |
| HO1 | $F_{2,6} = 174$<br>$p < 0.0001$ | ND | **0.01** | **0.0006** |

Differences to Nrf2 activation resulting from the level of oxygen in cell culture assays have been anecdotally reported. We therefore concurrently performed the overexpression experiments in HEK293 cells under conditions with 5% CO_2_ and ∼20% O_2_. Direct comparison of each transfection condition for each antibody indicated no significant difference between immunoreactivity of any condition (by two-way *t*-test without post-hoc correction). However, analysis of 20% O_2_ identified statistical significance for GCLC that had not been observed at 5% O_2_ (*F*_5,32_ = 7.1, *p*=0.003 by one-way ANOVA).

### Domain analysis of Bach1 and Bach2

Due to the fundamental differences between the inhibitory properties of Bach1 and Bach2, we characterized the major domains of Bach1 or Bach2. Previous studies suggest that the BTB binding domain would promote nuclear localization of Bach1 (Kanezaki et al., 2001; Yoshida et al., 1999), and the cytoplasmic localization signal (CLS) at the C-terminus of both Bach1 and Bach2 would promote nuclear export (Hoshino et al., 2000). However, others found that removal of the BTB or CLS domains promotes nuclear location of Bach1 (Suzuki et al., 2003). We therefore aimed to identify potential effects of removal of the BTB or CLS domains on Bach1 or Bach2-mediated transcriptional inhibition.

Removal of the CLS domain of Bach1 significantly decreased NQO1 and HO1 expression, when compared to wild-type Bach1 (**Fig 5**, Table 4). The same truncations did not significantly alter expression of other downstream antioxidant enzymes for either Bach1 or Bach2. Removal of the BTB domain of Bach1 trended to reduce NQO1 expression, however did not reach statistical significance when controlling for multiple comparisons. Removal of the DNA-binding domain (CNC-bZIP) completely abrogated the inhibitory activities of Bach1 and Bach2, resulting in antioxidant enzyme expression the similar to Nrf2/ pCMV expressing cells (in the absence of Bach1 or Bach2).

**Figure 5.**
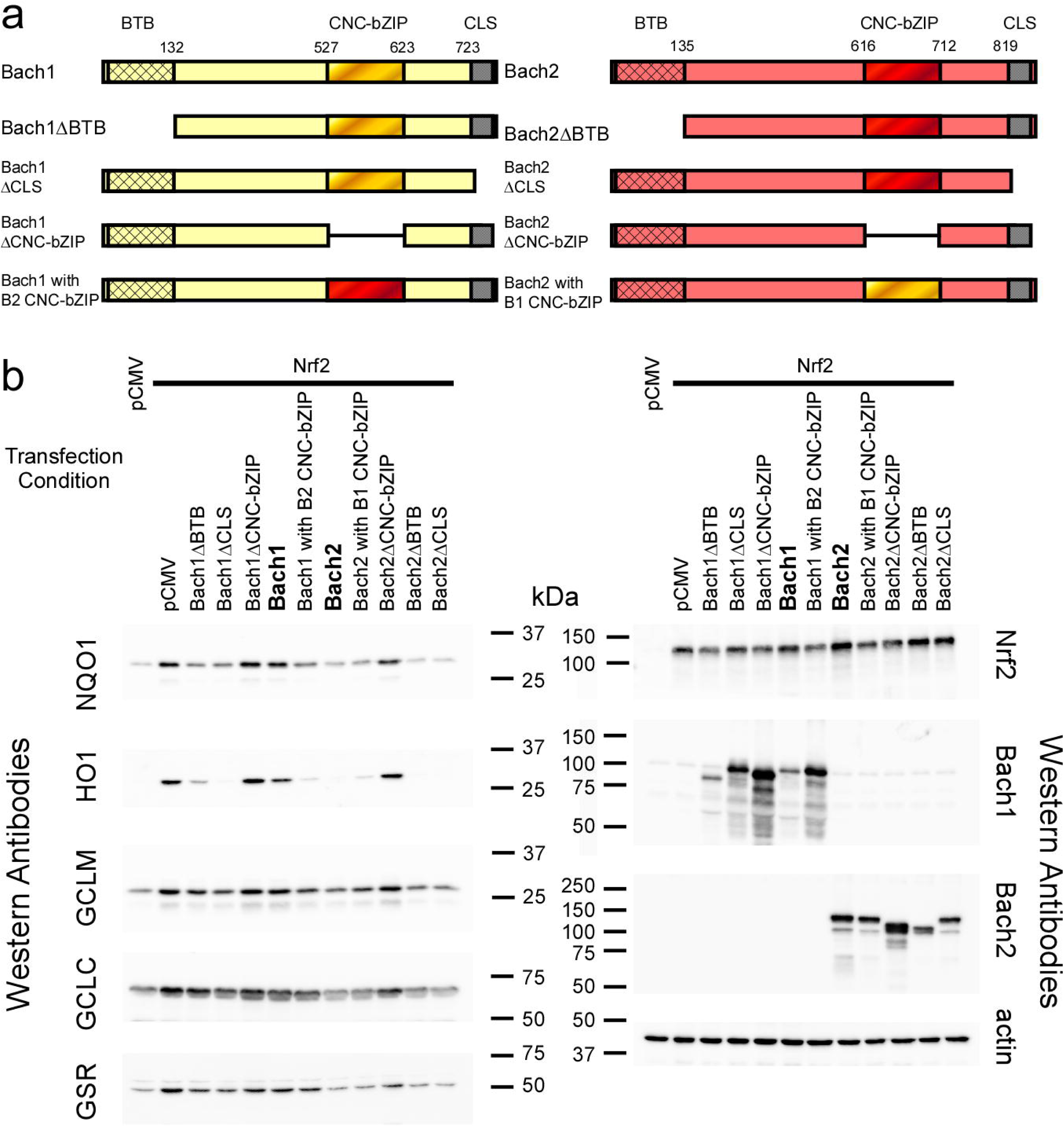
Bach1 and Bach2 deletion and hybrid mutations show relative functional domains. (a) Diagram depicting the following domains of Bach1 (depicted in yellow) and Bach2 (depicted in red): BTB (bric-a-brac, tramtrack, and broad complex; crosshatch lines), CNC-bZIP (cap’n collar basic leucine zipper; diagonal graduated-color lines), and CLS (cytoplasmic localization sequence; grey dots). The numbers provide directly above the domain edge demarcations are the amino acid residues at which the functional domains were truncated or changed. Indicated deletion and hybrid constructs are tested in (b). The following deletion mutations were created: ΔBTB (N-terminal BTB domain deletion), ΔCLS (C-terminal CLS deletion), and ΔCNC-bZIP (deletion of the DNA-binding domain). Hybrid mutations were made such that Bach1’s DNA binding domain was replaced with Bach2’s, and vice versa (Bach1 with B2 CNC-bZIP and Bach2 with B1 CNC-bZIP). Additional information is provided in the ‘Materials and Methods.’ (b) Western blot analysis of HEK293 cells transfected with Nrf2 in combination with modified Bach1 or Bach2 constructs, depicted in (a). Western blot analyses were performed with antibodies specific to NQO1, HO1, GCLM, GCLC, GSR, Nrf2, Bach1, Bach2, and actin. Shown are representative immunoblots of the typical data from three independent experiments.

**Table 4.**
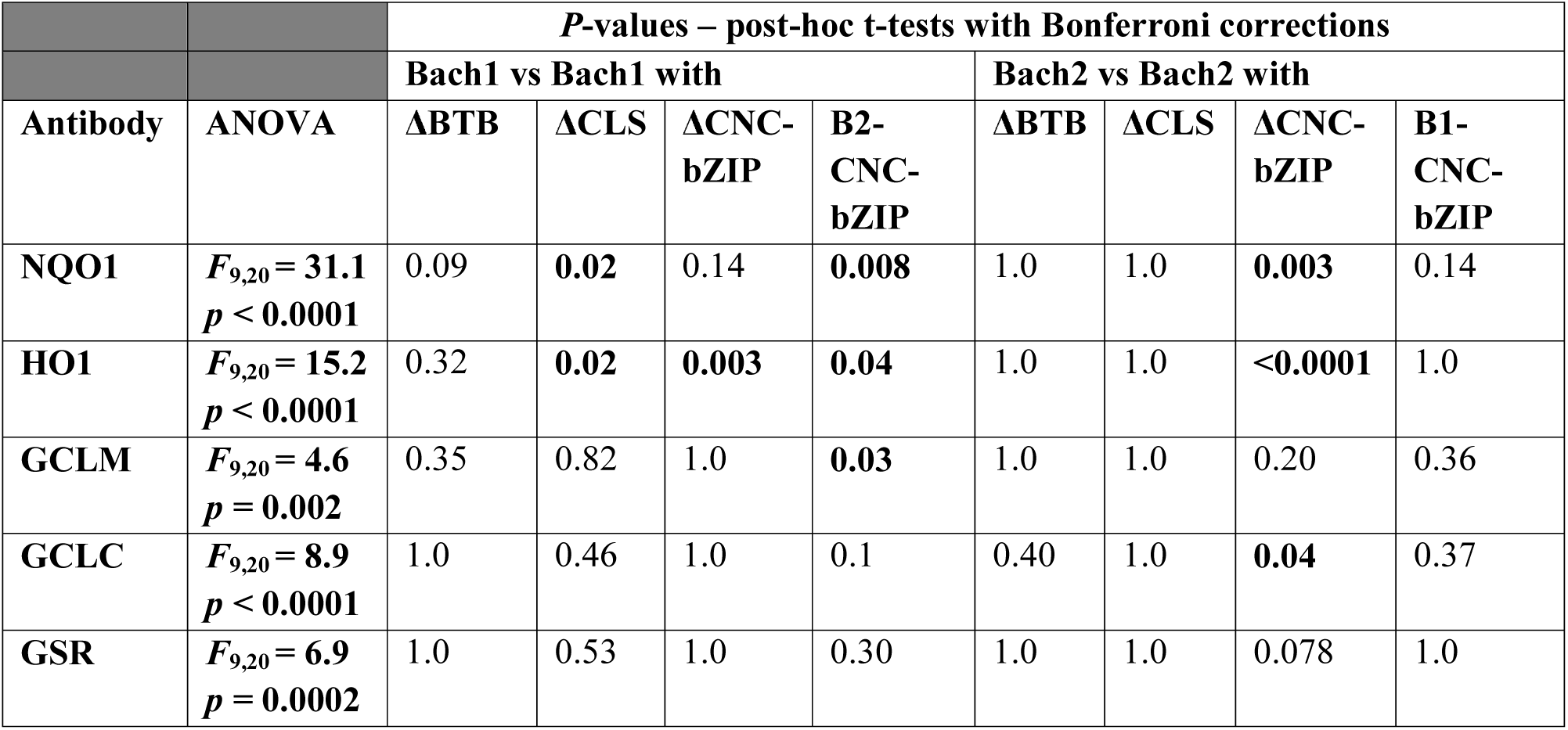
Statistical significance of differences between mutations of Bach1 and Bach2 cDNA transfections in HEK293 cells. One-way ANOVA tests were performed comparing all transfection conditions, followed by post-hoc two-way student t-tests with Bonferroni corrections (comparisons=5). pCMV and Nrf2/pCMV transfection conditions were omitted from analyses. Statistically significant *p*-values are shown in bold.

| Antibody | ANOVA | *P*-values – post-hoc t-tests with Bonferroni corrections |  |  |  |  |  |  |  |
| --- | --- | --- | --- | --- | --- | --- | --- | --- | --- |
|  |  | Bach1 vs Bach1 with |  |  |  | Bach2 vs Bach2 with |  |  |  |
|  |  | ΔBTB | ΔCLS | ΔCNC-bZIP | B2-CNC-bZIP | ΔBTB | ΔCLS | ΔCNC-bZIP | B1-CNC-bZIP |
| NQO1 | $F_{9,20} = 31.1$<br>$p < 0.0001$ | 0.09 | **0.02** | 0.14 | **0.008** | 1.0 | 1.0 | **0.003** | 0.14 |
| HO1 | $F_{9,20} = 15.2$<br>$p < 0.0001$ | 0.32 | **0.02** | **0.003** | **0.04** | 1.0 | 1.0 | **<0.0001** | 1.0 |
| GCLM | $F_{9,20} = 4.6$<br>$p = 0.002$ | 0.35 | 0.82 | 1.0 | **0.03** | 1.0 | 1.0 | 0.20 | 0.36 |
| GCLC | $F_{9,20} = 8.9$<br>$p < 0.0001$ | 1.0 | 0.46 | 1.0 | 0.1 | 0.40 | 1.0 | **0.04** | 0.37 |
| GSR | $F_{9,20} = 6.9$<br>$p = 0.0002$ | 1.0 | 0.53 | 1.0 | 0.30 | 1.0 | 1.0 | 0.078 | 1.0 |

To test the relative function of the DNA-binding domains in producing the observed Bach1 and Bach2 inhibitory activity, we created hybrid proteins of either Bach1 containing the Bach2 CNC-bZIP (Bach1 with B2 CNC-bZIP) or Bach2 containing the Bach1 CNC-bZIP (Bach2 with B1 CNC-bZIP). The Bach1-B2CNC-bZIP hybrid protein significantly decreased antioxidant enzyme expression when compared to wild-type Bach1, and more similar to that observed with Bach2 (**Fig 6**). Bach2-B1 CNC-bZIP hybrid expression did not significantly alter antioxidant enzyme expression when compared to wild-type Bach2. The resulting two hybrid proteins overall exhibited activities in between that of wild-type Bach1 and Bach2 proteins, suggesting that the DNA-binding domains of Bach1 and Bach2 only partially account for the differences between the proteins’ abilities to inhibit Nrf2 function.

**Figure 6.**
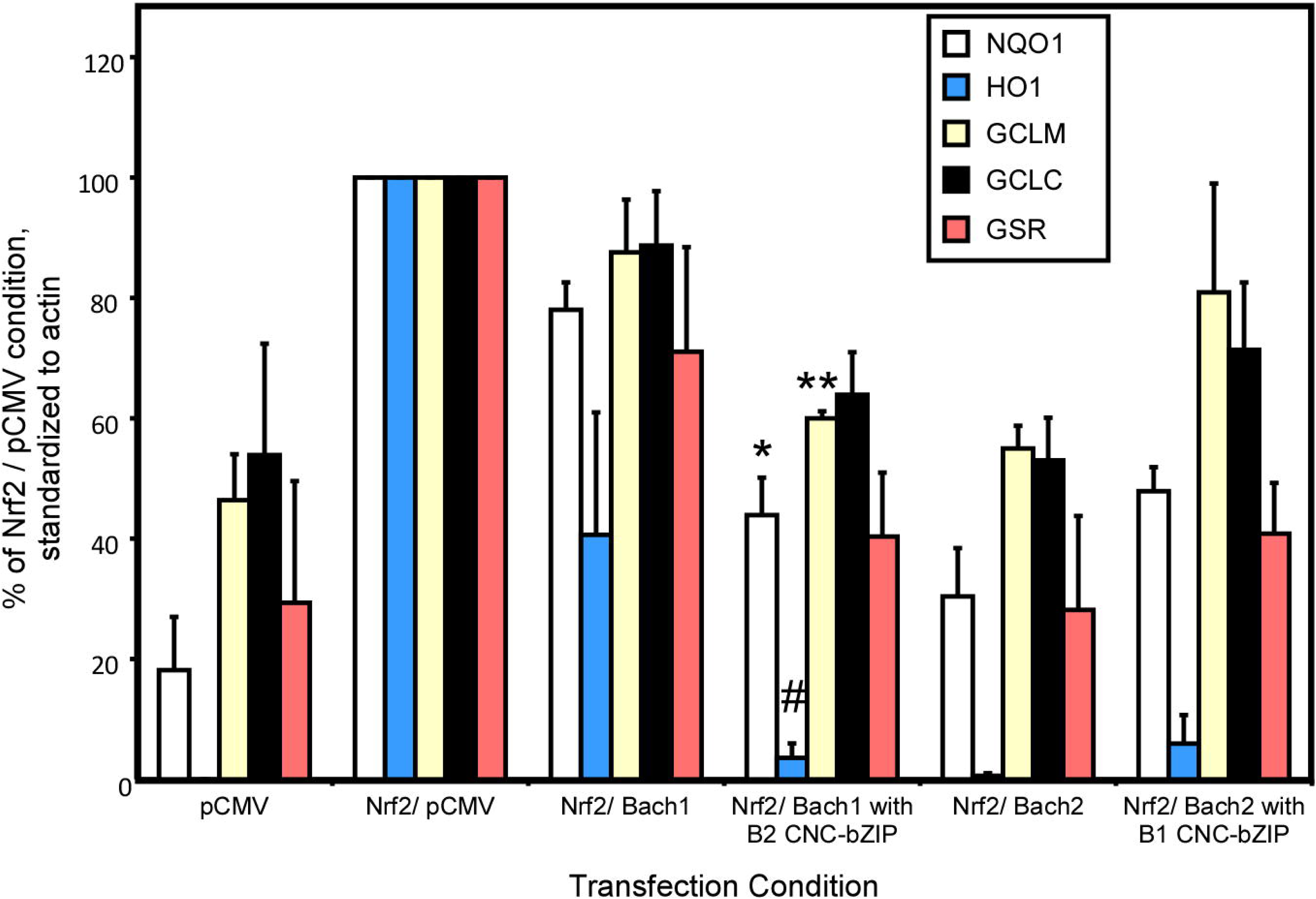
Differences in DNA-binding domains of Bach1 and Bach2 affect the inhibitory abilities of Bach1 and Bach2. Quantitative densitometry of Western blot data of HEK293 cells transfected with Nrf2 in combination with unmodified or hybrid mutations of Bach1 or Bach2 (Bach1 with B2 CNC-bZIP and Bach2 with B1 CNC-bZIP, respectively). Western blot analyses were performed with antibodies specific to NQO1, HO1, GCLM, GCLC, or GSR. Antibody reactivity was standardized to actin and provided as a percent of the ratio in Nrf2/ pCMV transfected cells. Statistical analyses were performed, comparing the hybrid Bach1 or Bach2 to the unmodified Bach1 or Bach2, respectively. Replacing the Bach1 CNC-bZIP domain with that of Bach2 significantly increased the inhibitory ability of Bach1 for for NQO1, HO1, and GCLM (*, *p*=0.0008; #, *p*=0.04; **, *p*=0.03, respectively). Data represent average ± SD for three independent experiments.

**Figure 7.**
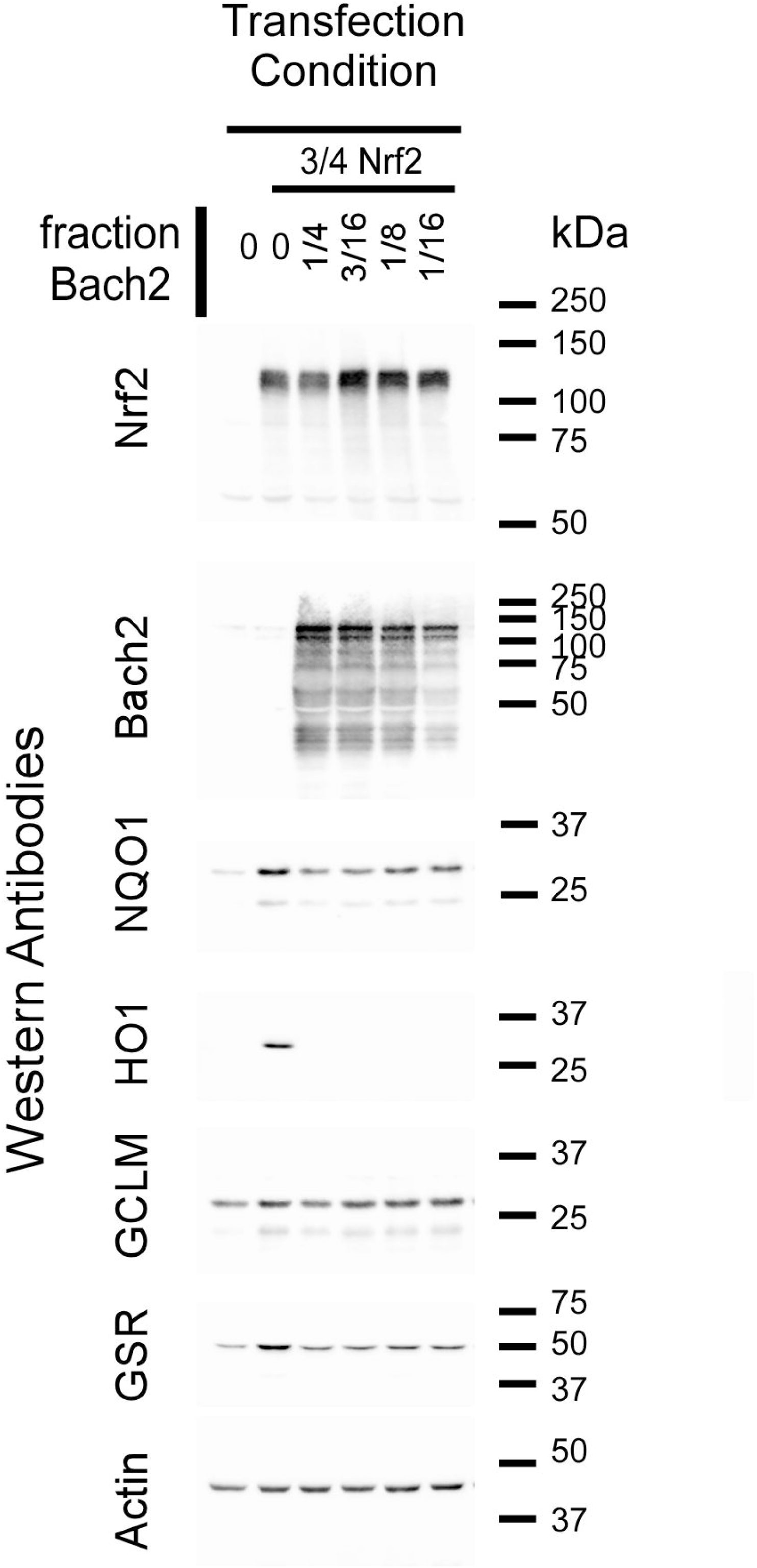
Relative Nrf2-mediated antioxidant enzyme expression with reduced Bach2 expression. Western blot analysis of HEK293 cells transiently transfected with a consistent amount of Nrf2 cDNA (3/4 of the transfection solution) and reducing amounts of Bach2 cDNA. The fraction of Bach2 cDNA ranged from 1/4 to 1/16 and was compared against 0 (Nrf2/ pCMV only). The transfection mix was buffered with pCMV vector to provide constant DNA amounts in each transfection condition. With the lowest transfection amount of Bach2 (1/16), significant inhibitory abilities of Bach2 remained. NQO1, GCLM, and GSR expression increased marginally and HO1 expression did not recover with reduced Bach2 expression. Shown are representative immunoblots of the typical data from three independent experiments.

### Inhibitory effects with reduced Bach2 expression

Trend differences in expression levels of Bach1 and Bach2 were observed with different mutations/ hybrid proteins, correlating with alterations in antioxidant enzyme expression levels. This observation suggests that expression levels could explain the differences in relative inhibitory activities between Bach1 and Bach2. We therefore reduced Bach2 expression to test a potential limit to Bach2’s inhibitory abilities. We titrated out Bach2 cDNA from the transfection solution mix, keeping Nrf2 cDNA at a steady level of 3/4 of the DNA and reducing Bach2 from 1/4 down to 1/16 of the DNA mix and buffering the remaining amount with pCMV vector. We then immunoblotted with antibodies specific to NQO1, HO1, GCLM, and additionally added glutathione-*S*-reductase (GSR), another antioxidant enzyme under ARE transcriptional control (**Fig 6**).

As previously observed, Nrf2 expression in the absence of Bach2 significantly increased the expression of all antioxidant enzymes, and co-expression of Bach2 reduced antioxidant enzyme expression to near basal levels with 1/4 of Bach2 cDNA. At the lowest transfection level of Bach2 tested (1/16 of DNA), significant inhibitory ability of Bach2 remained. The most dramatic was that of HO1, which did not recover with greatly diminished Bach2 expression. However, some recovery of expression of other antioxidants was observed (NQO1 and GCLM). This result supported the robust inhibitory abilities of Bach2 preventing Nrf2-mediated increases in antioxidant enzyme production that may be partially dependent on Bach2 expression levels.

### Effectiveness of chemical inhibitors against Bach1 and Bach2

Metals, such as cadmium chloride (CdCl_2_) and cobalt protoporphyrin (CoPP) can inhibit Bach1 and Bach2 function through nuclear export and/or degradation (Shan, Lambrecht, Donohue, & Bonkovsky, 2006; Suzuki et al., 2003). We examined the relative inhibitory properties of CdCl_2_ and CoPP by treating cells overexpressing Nrf2 with Bach1 or Bach2 with each of the inhibitors (**Fig 8**, **Fig 9**, Table 5, Table 6). Both CdCl_2_ and CoPP completely reversed Bach1’s inhibitory effect on HO1, consistent with previous studies (Shan et al., 2006; Suzuki et al., 2003). CdCl_2_ also partially reversed Bach2’s inhibitory effects on HO1, NQO1, and GCLM. CdCl_2_ and CoPP further promoted Nrf2 expression and activity in the absence of Bach overexpression, potentially through augmenting Nrf2 activation and/or inhibiting endogenous Bach1.

**Figure 8.**
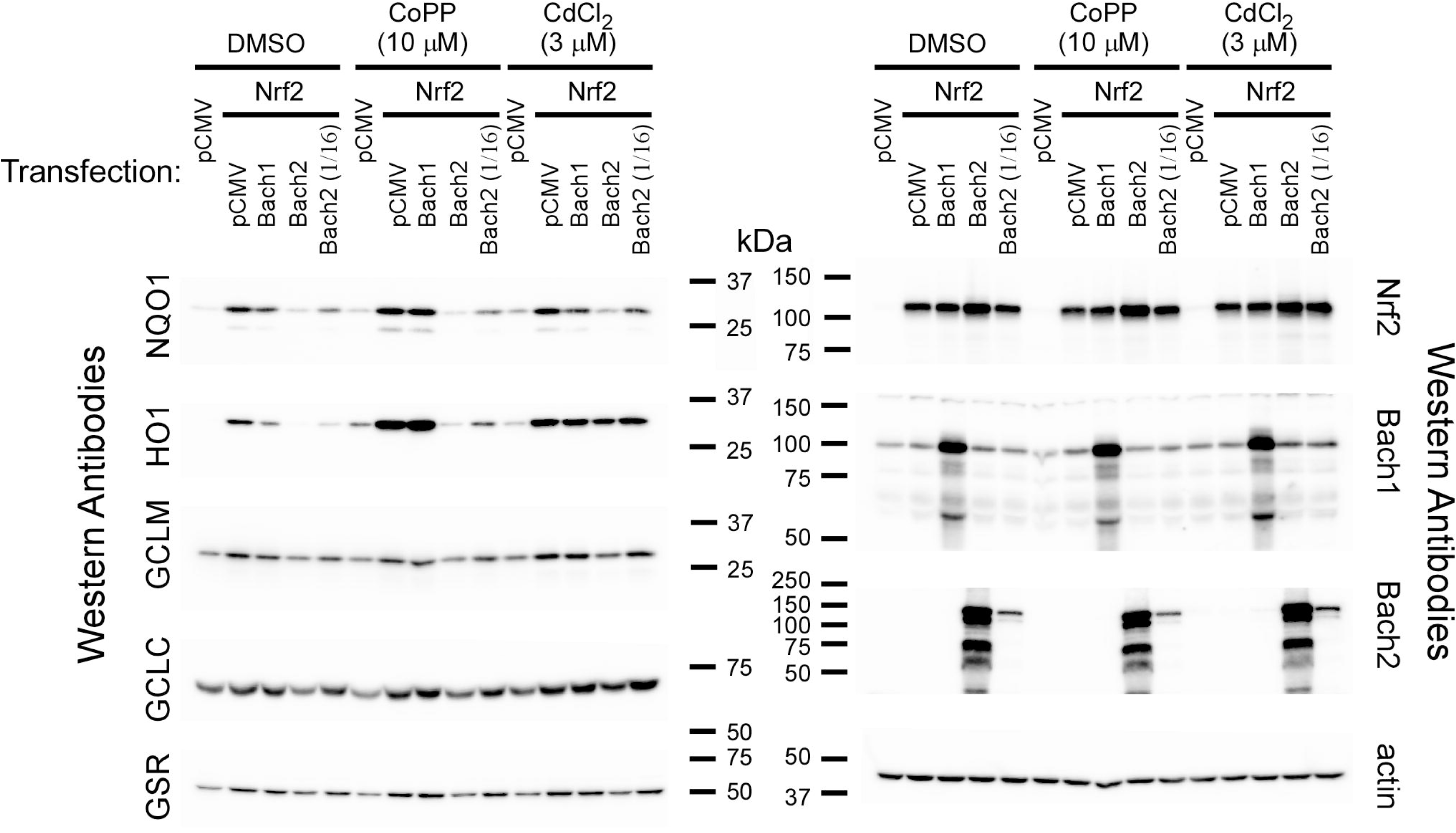
Bach inhibitors cobalt protoporphyrin (CoPP) and cadmium chloride (CdCl_2_) promote Nrf2-mediated antioxidant enzyme expression. Western blot analysis of HEK293 cells transiently transfected with a combination of Nrf2 cDNA with Bach1 or Bach2 cDNA and then treated with DMSO (vehicle control), CoPP (10 μM), or CdCl_2_ (3μM). Each cDNA represented 1/2 of the DNA in the transfection mix, except Nrf2/ Bach2 (1/16), where 1/2 of the cDNA was that of Nrf2, 7/16 was pCMV, and 1/16 was Bach2. Relative increases in antioxidant enzyme expression was observed when transfected cells were treated with CoPP or CdCl_2_ (see Figure 9). Shown are representative immunoblots of the typical data from three independent experiments.

**Figure 9.**
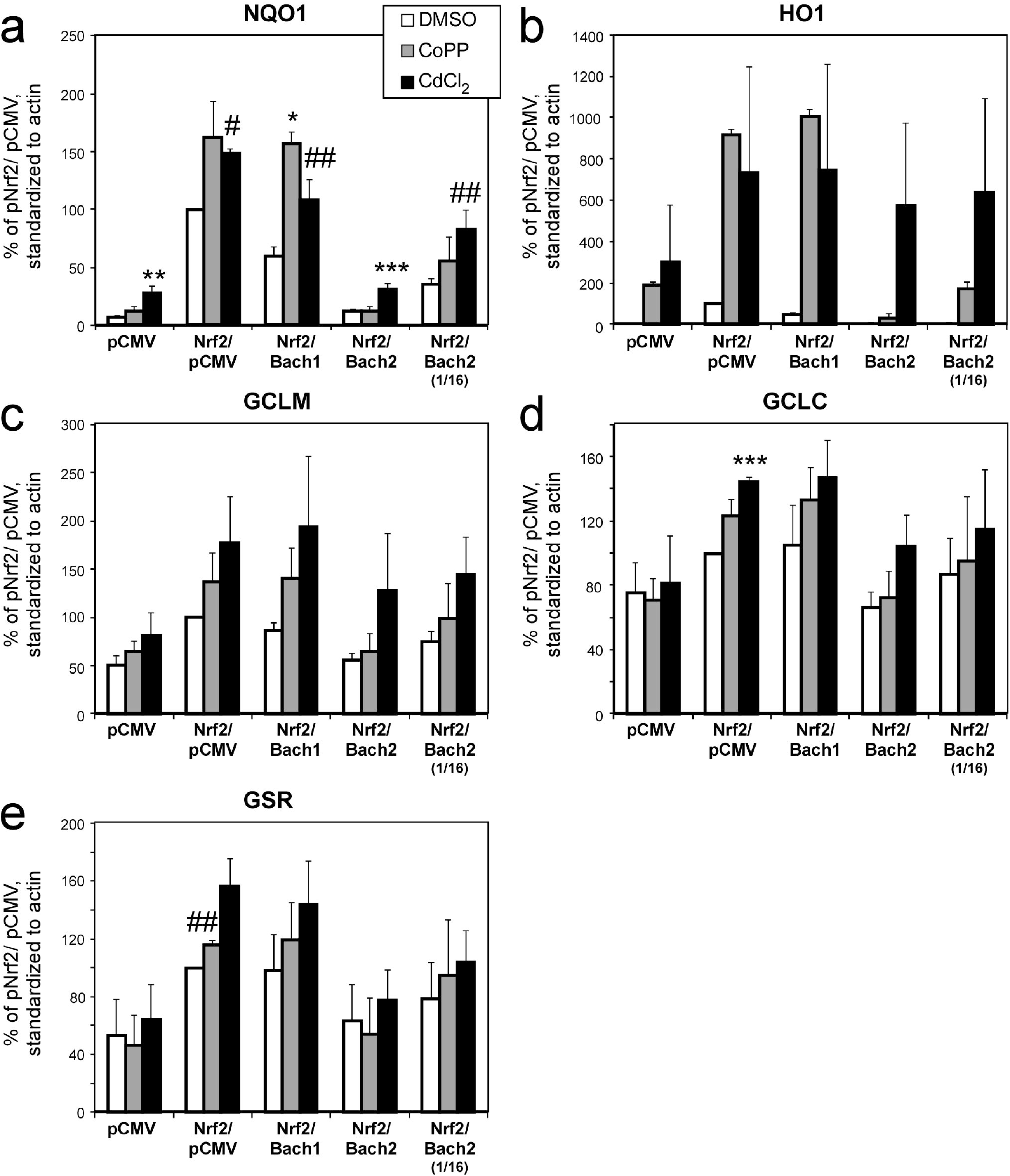
Quantitative analysis of cobalt protoporphyrin (CoPP) and cadmium chloride (CdCl_2_) treatment against Bach overexpressing HEK293 cells. Quantitative densitometry of Western blot data of antioxidant enzymes from HEK293 cells transfected with Nrf2 and pCMV, Bach1, Bach2, or 1/16 Bach2 and treated with DMSO, CoPP (10µM), or CdCl_2_ (3µM). Antioxidant enzymes of (a) NQO1, (b) HO1, (c) GCLM, (d) GCLC, and (e) GSR were assessed. Data was standardized to actin and quantitated as a percent of the Nrf2/pCMV transfection condition for each experiment. Both the transfection conditions and the drug treatments significantly affected expression levels of antioxidant enzymes (see Tables 4-5). Comparisons for CoPP or CdCl_2_ treatment were made against the DMSO treatment within each transfection condition (*, *p*=0.0004; **, *p*=0.006, #, *p*=0.004; ##, *p*=0.02, ***, *p*=0.003). Data represent average ± SD for three independent experiments.

**Table 5.** ANOVA analyses of Nrf2, Bach1, and Bach2 cDNA transfections in HEK293 cells with CoPP or CdCl_2_ treatment. Two-way ANOVA tests were performed comparing transfection conditions and drug treatments and interactions between the two factors (“Transfection * Drug”). Post-hoc two-way paired t-tests were performed comparing DMSO (control) treatment versus CoPP or CdCl_2_ to determine pairwise effects of each drug in altering protein expression as determined by each antibody. Statistically significant *p*-values are shown in bold.

| Antibody | ANOVA<br>Transfection | ANOVA<br>Drug treatment | ANOVA<br>Transfection *<br>Drug | *P*-values – paired t-tests |  |
| --- | --- | --- | --- | --- | --- |
|  |  |  |  | CoPP | CdCl<sub>2</sub> |
| NQO1 | $F_{4,30} = 177$<br>$p < 0.0001$ | $F_{2,30} = 45.8$<br>$p < 0.0001$ | $F_{8,30} = 9.99$<br>$p < 0.0001$ | **0.004** | **<0.0001** |
| HO1 | $F_{4,30} = 3.27$<br>$p = 0.02$ | $F_{2,30} = 10.7$<br>$p = 0.0003$ | $F_{8,30} = 1.39$<br>$p = 0.24$ | **0.007** | **<0.0001** |
| GCLM | $F_{4,30} = 8.57$<br>$p < 0.0001$ | $F_{2,30} = 16.8$<br>$p < 0.0001$ | $F_{8,30} = 0.599$<br>$p = 0.77$ | **0.001** | **<0.0001** |
| GCLC | $F_{4,30} = 10.7$<br>$p < 0.0001$ | $F_{2,30} = 8.0$<br>$p = 0.002$ | $F_{8,30} = 0.51$<br>$p = 0.84$ | 0.065 | **0.003** |
| GSR | $F_{4,30} = 18.0$<br>$p < 0.0001$ | $F_{2,30} = 7.7$<br>$p = 0.002$ | $F_{8,30} = 0.74$<br>$p = 0.66$ | 0.24 | **0.0002** |

**Table 6.** Statistical significance of CoPP or CdCl_2_ treatment in HEK293 cells transfected with Nrf2, Bach1, and Bach2 cDNA. Post-hoc tests were performed from the ANOVA analyses shown in Table 5 by two-way t-tests between DMSO (control) versus CoPP or CdCl_2_ for each transfection condition and antibody. Comparisons in the Nrf2/pCMV transfection were performed by one-sample t-tests, and all other comparisons were performed by student t-tests, each followed by Bonferroni corrections (comparisons=2). Statistically significant *p*-values are shown in bold.

| Antibody | *P*-values – post-hoc t-tests with Bonferroni corrections |  |  |  |  |  |  |  |  |  |
| --- | --- | --- | --- | --- | --- | --- | --- | --- | --- | --- |
|  | pCMV |  | Nrf2/pCMV |  | Nrf2/Bach1 |  | Nrf2/Bach2 |  | Nrf2/ Bach2 1/16 |  |
|  | CoPP | CdCl<sub>2</sub> | CoPP | CdCl<sub>2</sub> | CoPP | CdCl<sub>2</sub> | CoPP | CdCl<sub>2</sub> | CoPP | CdCl<sub>2</sub> |
| NQO1 | 0.22 | **0.006** | 0.14 | **0.004** | **0.0004** | **0.02** | 1.0 | **0.003** | 0.34 | **0.02** |
| HO1 | 0.12 | 0.26 | 0.33 | 0.33 | 0.13 | 0.15 | 0.057 | 0.14 | 0.077 | 0.15 |
| GCLM | 0.39 | 0.21 | 0.33 | 0.22 | 0.09 | 0.13 | 0.93 | 0.21 | 0.68 | 0.08 |
| GCLC | 1.0 | 1.0 | 0.13 | **0.003** | 0.39 | 0.20 | 1.0 | 0.078 | 1.0 | 0.63 |
| GSR | 1.0 | 1.0 | **0.02** | 0.070 | 0.52 | 0.14 | 1.0 | 0.97 | 1.0 | 0.28 |

Although direct comparisons between drug treatments and DMSO control did not consistently show statistically significant differences, the data support inhibitory effects of CoPP on Bach1 and inhibitory effects of CdCl_2_ on Bach1 and Bach2, and potential increased Nrf2 activation with both drugs, with statistical significance potentially limited by experimental variability and conservative multiple comparison testing.

### Bach1 and Bach2 expression in dopaminergic neuroblastoma cells

We aimed to extend our analysis of Bach1 and Bach2 to a dopaminergic model cell line, which could more accurately represent the potential inhibitory activities of Bach1 and Bach2 *in vivo*. We therefore examined expression levels in two human-derived neuroblastoma cell lines (IMR-32 and LUHMES), each of which have a dopaminergic phenotype after differentiation (Gupta, Notter, Felten, & Gash, 1985; Koutsilieri et al., 1996; Lotharius et al., 2005; Richards & Sadee, 1986; Scholz et al., 2011).

Under basal conditions, Bach1 was modestly expressed in all of the cell lines examined, and Bach2 expression was barely detectable (**Fig 10**). Interestingly, after differentiation of the two different neuroblastoma lines, Bach1 expression was reduced, and a lower molecular weight band (at ∼ 70kDa) appeared with the anti-Bach1 antibody. Bach2 expression, however, increased after neuronal differentiation, consistent with previous reports of differentiation of other neuronal cell lines (Hoshino & Igarashi, 2002; Shim, Rosner, Freilinger, Lubec, & Hengstschlager, 2006). No significant alterations in expression were observed for cells were maintained at 5% CO_2_ and 5% O_2_ versus 5% CO_2_ and ∼ 20% O_2_.

**Figure 10.**
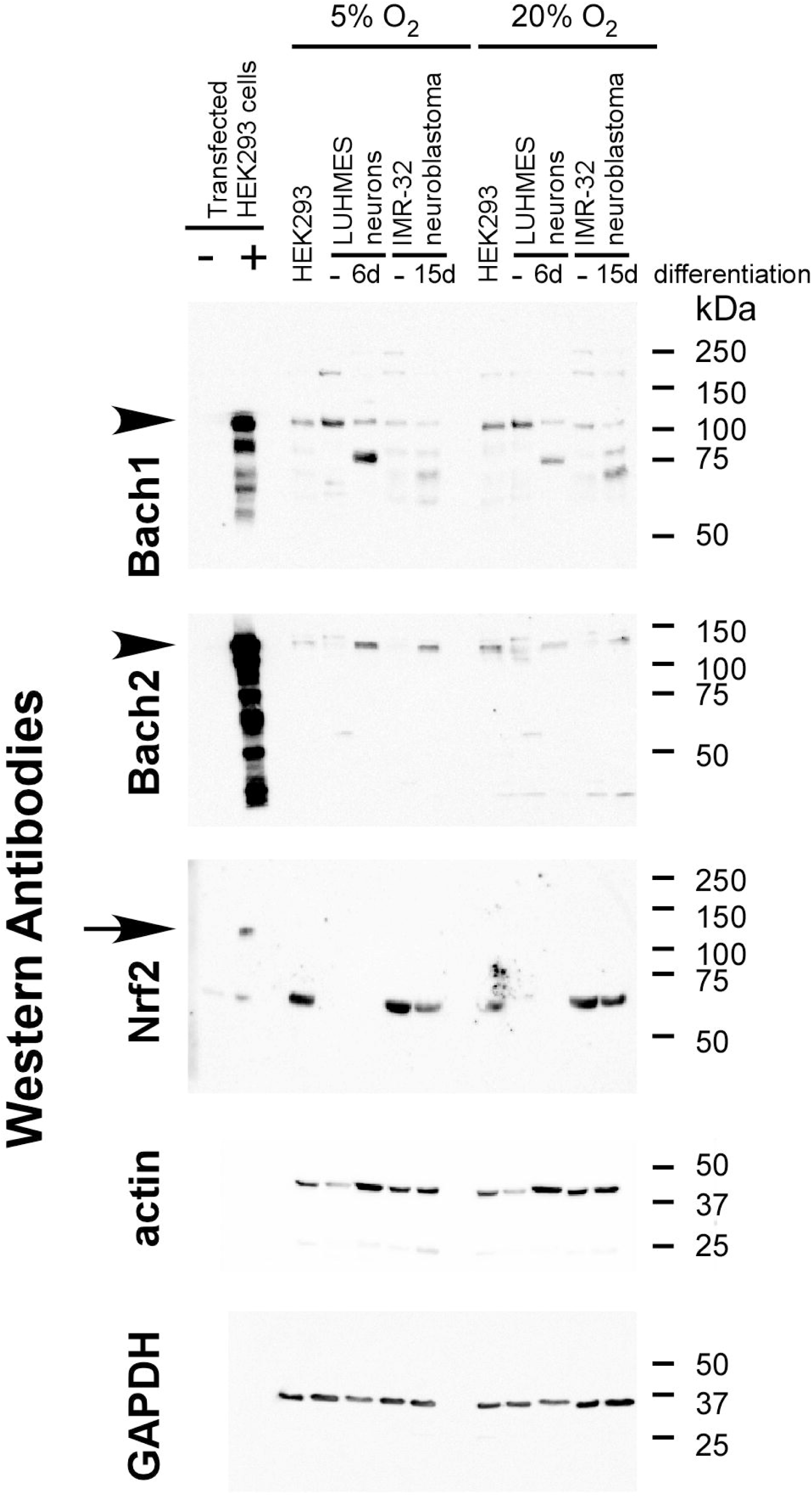
Expression profiles of endogenous Bach1, Bach2, and Nrf2 in dopaminergic neuronal cell lines. Representative Western blots of endogenously expressed Nrf2, Bach1, and Bach2 in HEK293 cells, LUHMES neurons, and IMR-32 neuroblastoma maintained under 5% O_2_ or ∼20% O_2_. As a comparator for molecular weight, transfected HEK293 cells were used as for each protein (“-“ for pCMV control and “+” for overexpressed proteins). 7.5μg protein of – and + controls were assessed concurrently with 30μg protein of each cell sample. LUHMES and IMR-32 cells were also differentiated for 6 days (6d) or 15 days (15d), respectively. Data show that after differentiation, both LUHMES and IMR-32 cells increased in Bach2 expression and decreased in full-length Bach1 expression. Neuronal lines also exhibited increased expression of a truncated forms of Bach1 after differentiation. Nrf2 expression is not readily observed at the upper molecular weight form consistent with the positive control in HEK293 cells (∼120kDa; arrowhead).

### Nrf2 activation and Bach inhibition in dopaminergic neuroblastoma

We utilized IMR-32 neuroblastoma cells to characterize the inhibitory properties of Bach1 and Bach2 on Nrf2-mediated antioxidant enzyme expression, taking advantage of the change in expressions between Bach1 and Bach2 with differentiation. Monomethyl fumarate (MMF) is an *in vitro* Nrf2 activator and the active metabolite of dimethyl fumarate (Linker et al., 2011; Scannevin et al., 2012). We therefore used MMF to activate Nrf2-dependent expression of antioxidant enzymes NQO1, HO1, GCLM, GCLC, and GSR (**Figs 11-12**). MMF treatment on its own, and concomitant treatment with CoPP or CdCl_2_ treatment promoted Nrf2 activity and antioxidant enzyme expression. These increases in many cases increased that of Nrf2 expression. In addition, Bach1 expression increased with Nrf2 activation. However, a stark difference was observed after neuronal differentiation. Differentiation blunted MMF, CoPP, and CdCl_2_-induced increases in NQO1, HO1, GCLM, GSR, and Nrf2 expression. These changes correlated with reduced Bach1 – full length expression and increased Bach2 and Bach1 – truncated expression.

**Figure 11.**
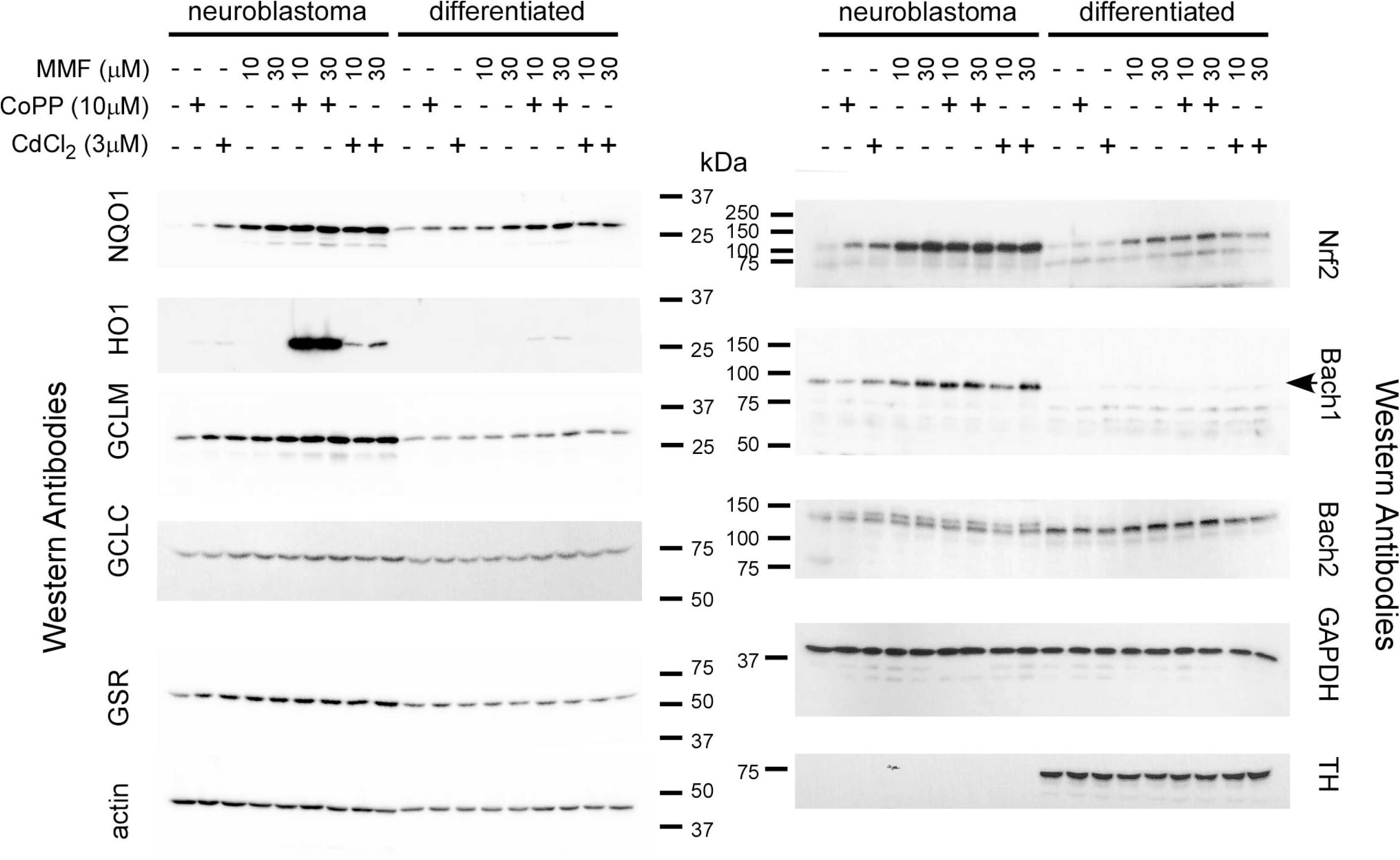
Differentiation of IMR-32 cells blunts Nrf2-mediated antioxidant enzyme expression. Western blot analysis of IMR-32 neuroblastoma treated with the Nrf2 activator monomethyl fumarate (MMF) with or without the Bach inhibitors cobalt protoporphyrin (CoPP) or cadmium chloride (CdCl_2_). Nrf2 activation, as shown by increases in Nrf2 expression and antioxidant enzyme expression, was observed when cells were in their neuroblastoma form. After 16d of differentiation, full length Bach1 expression (arrow) was diminished and Bach2 expression was increased, corresponding with reduced MMF-induced responses. Addition of CoPP orCdCl_2_ only minimally promoted Nrf2 responses in differentiated IMR-32 cells (See Figure 12). Shown are representative immunoblots from three independent experiments.

**Figure 12.**
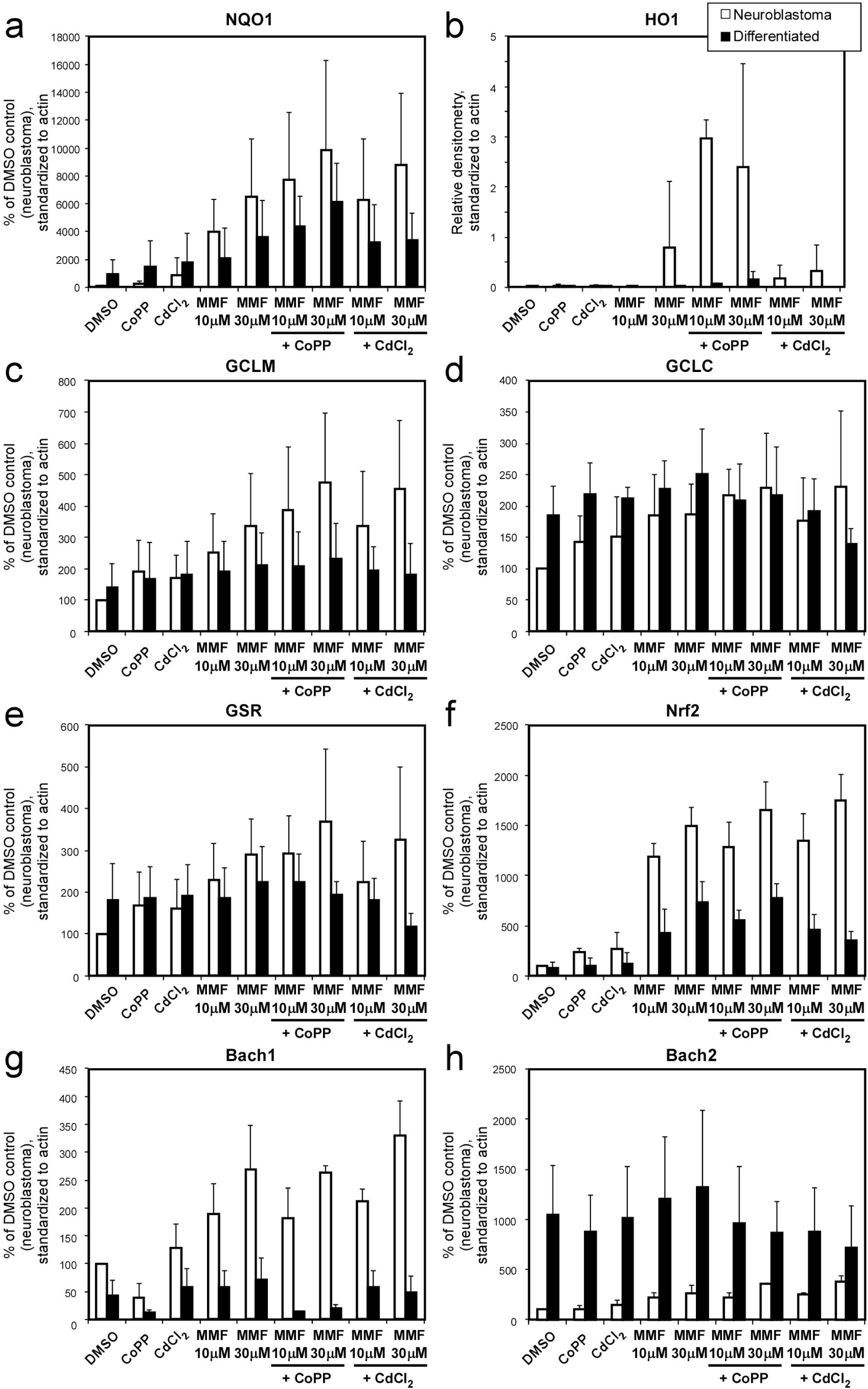
Quantitative analysis of IMR-32 cells with MMF, cobalt, and cadmium treatments in the absence/ presence of differentiation. Quantitative densitometry of Western blot data performed on IMR-32 cells in the neuroblastoma state (white) versus differentiated for 16 days (black) for antioxidant enzymes NQO1(a), HO1 (b), GCLM (c), GCLC (d), GSR (e), Nrf2 (f), Bach1 – full length (g), and Bach2 (h). Data was standardized to actin and then analyzed as a percent of DMSO control, except for HO1, where only relative ratio densitometry was analyzed due to the lack of immunoreactivity under control conditions. Overall trends of reduced Nrf2 responses after differentiation can be observed for NQO1, HO1, GCLM, and GSR. CoPP and CdCl_2_ increased some antioxidant responses prior to differentiation, but had a minimal response after differentiation (See Figure 13). Data represent average ± SD for 3 independent experiments.

Using the four variables of differentiation, MMF treatment, CoPP treatment, and CdCl_2_ treatment, and the interaction between these variables in a multi-factor ANOVA analysis, the relative contributions of each variable or interaction of variables were assessed for each antibody immunoreactivity (**Fig 13**; Table 7). Differentiation and MMF treatment contributed the most to the levels of observed antibody immunoreactivity, as did the interaction between these two variables. However, CoPP only significantly increased HO1 and decreased Bach1 (full-length), and CdCl_2_ did not significantly contribute to immunoreactivity levels. A post-hoc comparison between neuroblastoma and differentiated cells of the same treatments identified statistically significant effects for CoPP + MMF (10µM) for HO1, CdCl_2_ + MMF (10µM) for Bach1, and CdCl_2_ + MMF (30µM) for Nrf2 (*p*=0.001, *p*=0.009, *p*=0.009, respectively, by two-way t-test followed by Bonferroni corrections; Table 8).

**Figure 13.**
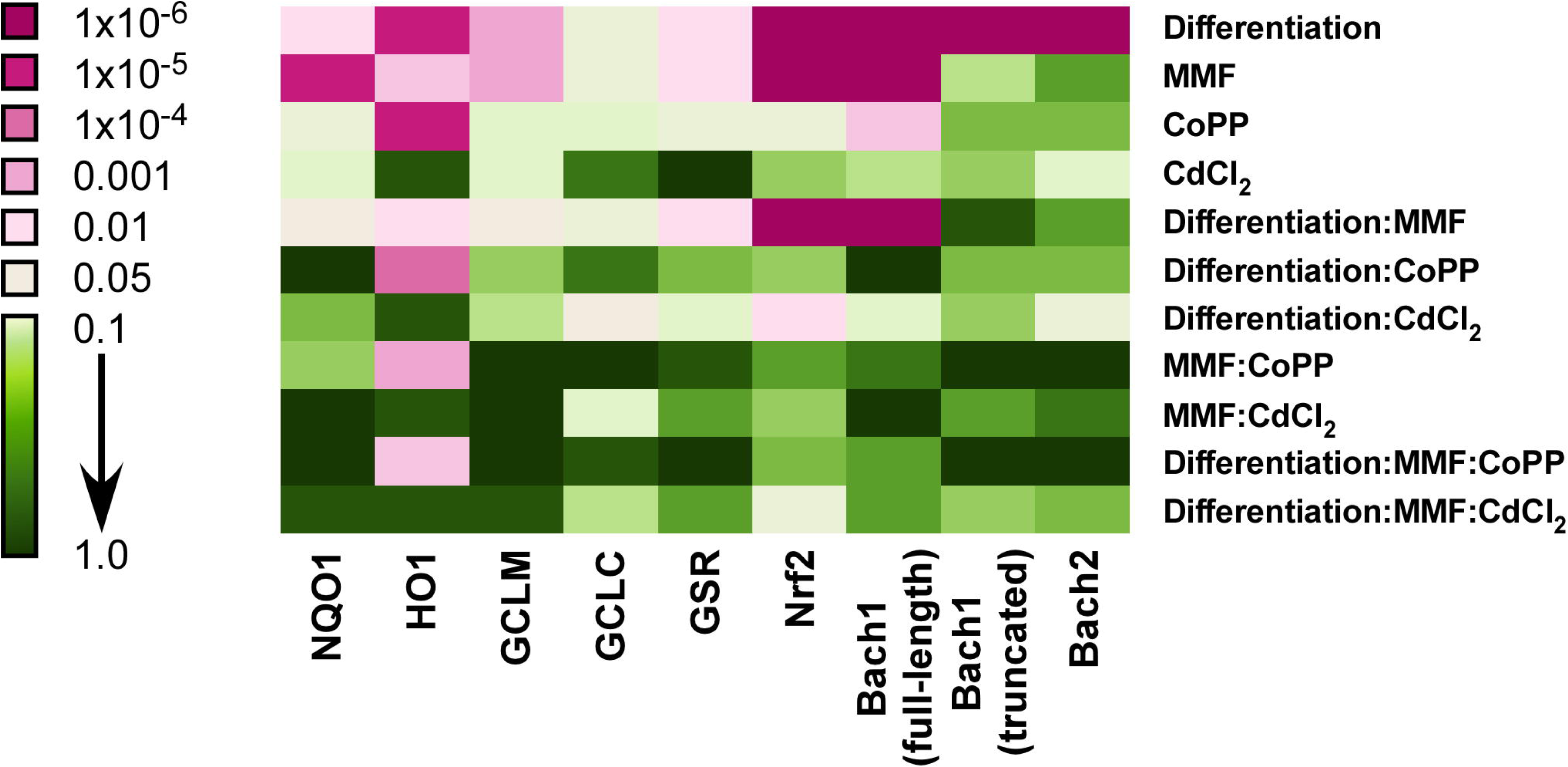
Variables associated with antioxidant enzyme response in IMR-32 cells. Multiple factor ANOVA analyses were performed for each antioxidant enzyme, Nrf2, Bach1 (full-length), Bach1 (truncated), and Bach2 with the variables of differentiation, MMF treatment, CoPP treatment, and CdCl_2_ treatment analyzed. Interactions between variables were additionally assessed (as represented as “:” between variable names). Represented is a colorimetric visualization of *p*-values based on the resulting *F*-statistics for each variable or interaction, analyzed for each antibody’s standardized quantitative densitometry (antibodies indicated on the x-axis) for data shown in Figure 12. Pink coloring represents statistical significance (*p* <0.05).

**Table 7.** ANOVA analyses of IMR-32 cells with treatment conditions MMF, CoPP, and/or CdCl_2_ in the absence or presence of differentiation. Two-way ANOVA tests were performed comparing Differentiation (Diff), MMF treatment, CoPP treatment, or CdCl_2_ treatment as a multi-factor analysis. In addition, the interactions between factors were assessed (indicated by “*” in the heading). Statistically significant *p*-values are shown in bold.

| Antibody | Multi-factor ANOVA analyses |  |  |  |  |  |  |  |  |  |  |
| --- | --- | --- | --- | --- | --- | --- | --- | --- | --- | --- | --- |
|  | Diff | MMF | CoPP | CdCl<sub>2</sub> | Diff *<br>MMF | Diff *<br>CoPP | Diff *<br>CdCl<sub>2</sub> | MMF *<br>CoPP | MMF<br>*<br>CdCl<sub>2</sub> | Diff *<br>MMF *<br>CoPP | Diff *<br>MMF<br>*<br>CdCl<sub>2</sub> |
| **NQO1** | $F_{1,36} = 5.03$<br>$p=0.03$ | $F_{2,36} = 14.2$<br>$p<0.0001$ | $F_{1,36} = 2.8$<br>$p=0.10$ | $F_{1,36} = 1.3$<br>$p=0.27$ | $F_{2,36} = 3.14$<br>$p=0.055$ | $F_{1,36} = 0.001$<br>$p=0.98$ | $F_{1,36} = 0.33$<br>$p=0.57$ | $F_{2,36} = 0.75$<br>$p=0.48$ | $F_{2,36} = 0.07$<br>$p=0.94$ | $F_{2,36} = 0.05$<br>$p=0.95$ | $F_{2,36} = 0.13$<br>$p=0.88$ |
| **HO1** | $F_{1,36} = 7.0$<br>$p<0.0001$ | $F_{2,36} = 1.9$<br>$p=0.009$ | $F_{1,36} = 8.2$<br>$p<0.0001$ | $F_{1,36} = 0.02$<br>$p=0.81$ | $F_{2,36} = 1.7$<br>$p=0.02$ | $F_{1,36} = 6.81$<br>$p=0.0001$ | $F_{1,36} = 0.02$<br>$p=0.82$ | $F_{2,36} = 2.24$<br>$p=0.005$ | $F_{2,36} = 0.08$<br>$p=0.80$ | $F_{2,36} = 1.98$<br>$p=0.008$ | $F_{2,36} = 0.08$<br>$p=0.80$ |
| **GCLM** | $F_{1,36} = 9.49$<br>$p=0.004$ | $F_{2,36} = 6.54$<br>$p=0.004$ | $F_{1,36} = 1.13$<br>$p=0.30$ | $F_{1,36} = 1.3$<br>$p=0.27$ | $F_{2,36} = 3.22$<br>$p=0.052$ | $F_{1,36} = 0.53$<br>$p=0.47$ | $F_{1,36} = 0.92$<br>$p=0.34$ | $F_{2,36} = 0.05$<br>$p=0.96$ | $F_{2,36} = 0.008$<br>$p=0.99$ | $F_{2,36} = 0.02$<br>$p=0.98$ | $F_{2,36} = 0.15$<br>$p=0.87$ |
| **GCLC** | $F_{1,36} = 2.48$<br>$p=0.12$ | $F_{2,36} = 2.31$<br>$p=0.11$ | $F_{1,36} = 1.23$<br>$p=0.28$ | $F_{1,36} = 0.07$<br>$p=0.79$ | $F_{2,36} = 2.38$<br>$p=0.11$ | $F_{1,36} = 0.07$<br>$p=0.79$ | $F_{1,36} = 2.93$<br>$p=0.10$ | $F_{2,36} = 0.002$<br>$p=1.00$ | $F_{2,36} = 1.29$<br>$p=0.29$ | $F_{2,36} = 0.15$<br>$p=0.86$ | $F_{2,36} = 1.18$<br>$p=0.32$ |
| **GSR** | $F_{1,36} = 4.66$<br>$p=0.04$ | $F_{2,36} = 4.47$<br>$p=0.02$ | $F_{1,36} = 2.15$<br>$p=0.15$ | $F_{1,36} = 0.005$<br>$p=0.94$ | $F_{2,36} = 5.15$<br>$p=0.01$ | $F_{1,36} = 0.41$<br>$p=0.53$ | $F_{1,36} = 1.15$<br>$p=0.29$ | $F_{2,36} = 0.15$<br>$p=0.86$ | $F_{2,36} = 0.48$<br>$p=0.62$ | $F_{2,36} = 0.004$<br>$p=1.00$ | $F_{2,36} = 0.46$<br>$p=0.64$ |
| **Nrf2** | $F_{1,36} = 182$<br>$p<0.0001$ | $F_{2,36} = 155$<br>$p<0.0001$ | $F_{1,36} = 2.21$<br>$p=0.15$ | $F_{1,36} = 0.68$<br>$p=0.42$ | $F_{2,36} = 33.6$<br>$p<0.0001$ | $F_{1,36} = 0.63$<br>$p=0.43$ | $F_{1,36} = 6.39$<br>$p=0.02$ | $F_{2,36} = 0.39$<br>$p=0.68$ | $F_{2,36} = 0.90$<br>$p=0.42$ | $F_{2,36} = 0.57$<br>$p=0.57$ | $F_{2,36} = 2.1$<br>$p=0.14$ |
| **Bach1 – full length** | $F_{1,36} = 118$<br>$p<0.0001$ | $F_{2,36} = 19.3$<br>$p<0.0001$ | $F_{1,36} = 8.59$<br>$p=0.006$ | $F_{1,36} = 1.08$<br>$p=0.31$ | $F_{2,36} = 16.3$<br>$p<0.0001$ | $F_{1,36} = 0.006$<br>$p=0.94$ | $F_{1,36} = 1.45$<br>$p=0.24$ | $F_{2,36} = 0.28$<br>$p=0.76$ | $F_{2,36} = 0.033$<br>$p=0.97$ | $F_{2,36} = 0.30$<br>$p=0.67$ | $F_{2,36} = 0.46$<br>$p=0.63$ |
| **Bach1 – truncated** | $F_{1,36} = 54.3$<br>$p<0.0001$ | $F_{2,36} = 1.02$<br>$p=0.37$ | $F_{1,36} = 0.30$<br>$p=0.59$ | $F_{1,36} = 0.49$<br>$p=0.49$ | $F_{2,36} = 0.21$<br>$p=0.81$ | $F_{1,36} = 0.35$<br>$p=0.56$ | $F_{1,36} = 0.54$<br>$p=0.47$ | $F_{2,36} = 0.045$<br>$p=0.96$ | $F_{2,36} = 0.47$<br>$p=0.63$ | $F_{2,36} = 0.045$<br>$p=0.96$ | $F_{2,36} = 0.82$<br>$p=0.45$ |
| **Bach2** | $F_{1,36} = 58.7$<br>$p<0.0001$ | $F_{2,36} = 0.37$<br>$p=0.69$ | $F_{1,36} = 0.36$<br>$p=0.55$ | $F_{1,36} = 1.15$<br>$p=0.29$ | $F_{2,36} = 0.42$<br>$p=0.66$ | $F_{1,36} = 0.38$<br>$p=0.54$ | $F_{1,36} = 2.58$<br>$p=0.12$ | $F_{2,36} = 0.01$<br>$p=0.99$ | $F_{2,36} = 0.36$<br>$p=0.70$ | $F_{2,36} = 0.03$<br>$p=0.97$ | $F_{2,36} = 0.55$<br>$p=0.58$ |

**Table 8.** Statistical significance of differentiation of IMR-32 cells for each of the treatment conditions. Post-hoc tests were performed from the ANOVA analyses shown in Table 7 by two-way t-tests between IMR-32 cells for each treatment condition indicated comparing undifferentiated versus differentiated cells. Comparisons between DMSO treatment conditions were performed by one-sample t-tests, and all other comparisons were performed by student t-tests, each followed by Bonferroni corrections (comparisons=9). Statistically significant *p*-values are shown in bold.

|  | *P*-values – post-hoc t-tests with Bonferroni corrections |  |  |  |  |  |  |  |  |
| --- | --- | --- | --- | --- | --- | --- | --- | --- | --- |
| Antibody | DMSO | CoPP | CdCl<sub>2</sub> | MMF<br>(10μM) | MMF<br>(30μM) | MMF<br>(10μM) +<br>CoPP | MMF<br>(30μM)<br>+ CoPP | MMF<br>(10μM) +<br>CdCl<sub>2</sub> | MMF<br>(30μM) +<br>CdCl<sub>2</sub> |
| NQO1 | 0.25 | 1.0 | 1.0 | 1.0 | 1.0 | 1.0 | 1.0 | 1.0 | 1.0 |
| HO1 | ND | 1.0 | 1.0 | 1.0 | 1.0 | 1.0 | **0.001** | 1.0 | 1.0 |
| GCLM | 0.42 | 1.0 | 1.0 | 1.0 | 1.0 | 1.0 | 1.0 | 1.0 | 1.0 |
| GCLC | 0.12 | 1.0 | 1.0 | 1.0 | 1.0 | 1.0 | 1.0 | 1.0 | 1.0 |
| GSR | 0.22 | 1.0 | 1.0 | 1.0 | 1.0 | 1.0 | 1.0 | 1.0 | 1.0 |
| Nrf2 | 0.47 | 0.50 | 1.0 | 0.08 | 0.07 | 0.08 | 0.08 | 0.06 | **0.009** |
| Bach1-full | 0.11 | 0.28 | 0.21 | 0.057 | 0.11 | 0.47 | 0.12 | **0.009** | 0.11 |
| Bach1 –<br>truncated | 0.10 | 0.49 | 0.33 | 0.37 | 0.27 | 0.056 | 1.0 | 1.0 | 0.87 |
| Bach2 | 0.11 | 0.18 | 0.40 | 0.45 | 0.68 | 0.80 | 0.41 | 0.63 | 1.0 |

The overall results show a relative response of antioxidants to MMF, which was blunted or in some cases prevented by differentiation, and a response of HO1 to CoPP that was consistent with a reduction of Bach1 expression with CoPP treatment, but only in the neuroblastoma/ non-differentiated state.

### Bach1 and Bach2 interact through their BTB domains

Bach chemical inhibitors were largely ineffective in differentiated IMR-32 cells, which express Bach2 and predominantly truncated forms of Bach1. CdCl_2_, which was more effective against Bach1 in transfected HEK293 cells, depends on the C-terminal CLS domain (Suzuki et al., 2003). Additionally, CoPP, which was not very effective against Bach2 in transfected HEK293 cells, had a small (although not statistically significant) trend with some antibodies in differentiated IMR-32 cells. One potential explanation for the data could be a cooperative relationship between the truncated Bach1 and Bach2 expression in the differentiated IMR-32 cells. Truncated forms of Bach1 may facilitate nuclear localization of other Bach proteins through multimeric binding of the BTB domain (Kanezaki et al., 2001). We therefore tested for an interaction between Bach1 and Bach2 BTB domains.

Co-immunoprecipitations was performed with Bach2 against Bach1 or the BTB domains of Bach1 or Bach2 in transfected HEK293 cells, followed by Western blot analysis, to identify interactions between Bach1 and Bach2 through their BTB domains (**Fig 14**). Immunoprecipitation of Bach1, B1-BTB, and B2-BTB was observed with anti-Bach2, but not with normal rabbit IgG. Significant immunoprecipitation was observed despite a lack of significant depletion of Bach2 from the unbound lysate (**Fig 14b**). When all of the cDNAs were mixed for the immunoprecipitation (“mix all”, Fig 11D), B1-BTB and B2-BTB were precipitated in equal amounts. Myc-tagged B1-BTB and B2-BTB also successfully immunoprecipitated Bach2 with an anti-myc antibody. These data support interactions between Bach1 and Bach2 BTB domains for dimer or multimerization of Bach1 and Bach2 proteins.

**Figure 14.**
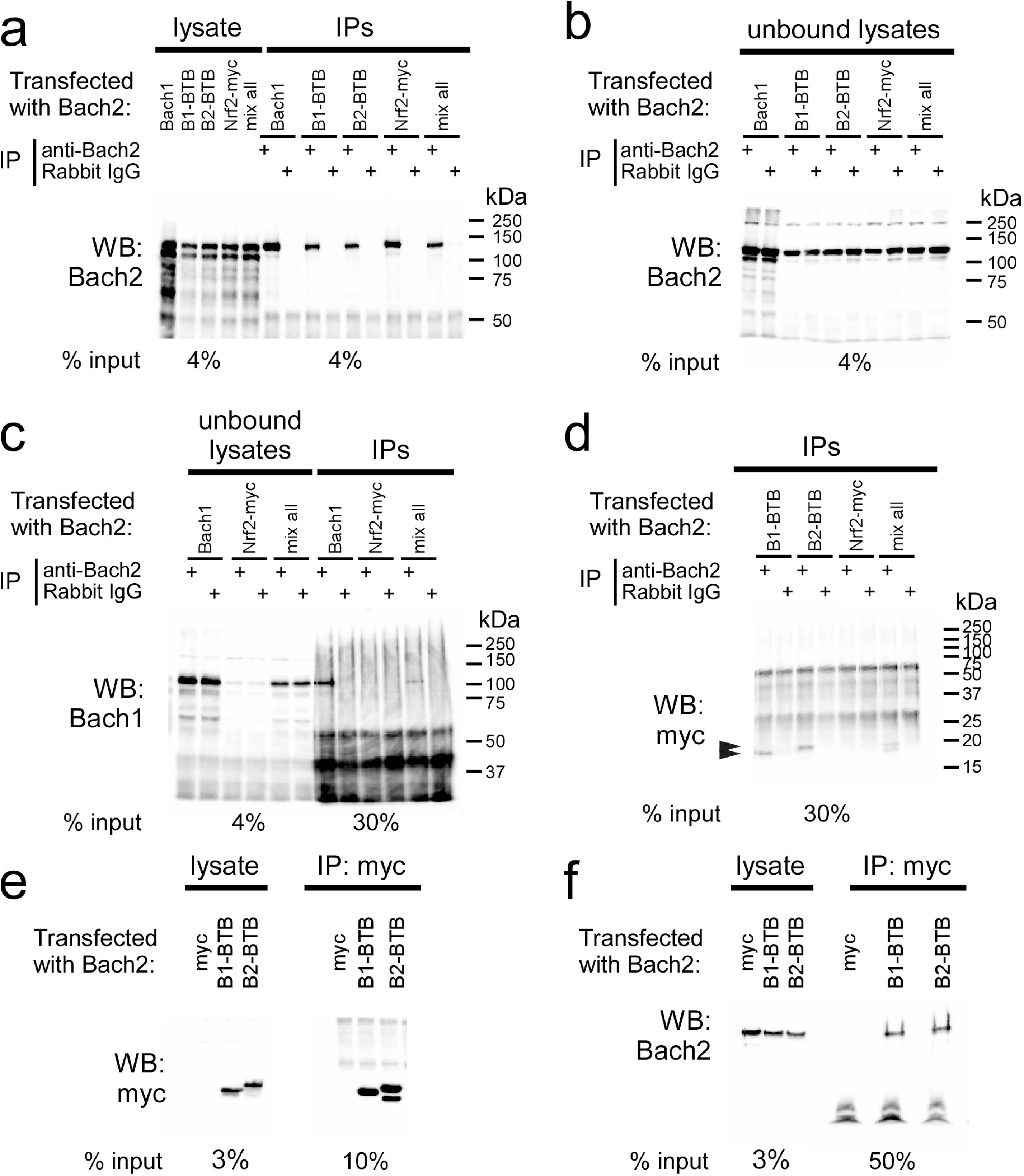
Interactions between Bach2 and Bach1 or BTB-myc performed by co-immunoprecipitation in transfected HEK293 cells. (a-d) HEK293 cells were co-transfected with Bach2 and either Bach1, Bach1-BTB-only-myc (B1-BTB), Bach2-BTB-only-myc (B2-BTB), Nrf2-myc, or a mix of all cDNAs in this experiment added in equal amounts (mix all), or (e,f) HEK293 cells were co-transfected with Bach2 and either B1-BTB, B2-BTB, or a myc control vector (myc). Immunoprecipitation (IP) was performed with antibodies anti-Bach2 or normal rabbit IgG (a-d) or anti-myc (e-f). Representative immunoblots show lysates (taken from the total protein pool), unbound lysates (taken from the remaining lysate after immunoprecipitation), or IP (taken from the proteins + antibodies bound to the beads). Each IP was performed with 250 µg of total cell lysate and the percent input (fraction of the lysate for the condition) is indicated underneath Western blot conditions. Western blots performed on IPs with anti-Bach2 (a,f), anti-Bach1 (c), or anti-myc (d-arrowheads, e) show successfully immunoprecipitation of each with anti-Bach2 IP or anti-myc IP, but not IP with the Rabbit IgG control or with a control myc plasmids. Even though significant depletion of Bach2 in the unbound lysate was not observed (b), immunoprecipitation procedures were sufficient to show significant IP of target proteins. Of note, when all cDNAs were included in the transfection mixture (mix all), both B1-BTB and B2-BTB were observed in the IP in approximately equal quantities (d).

## Discussion

The current study provides characterization and comparisons of Bach1 and Bach2 activity to identify their relative contributions in preventing the protective Nrf2-mediated antioxidant responses. This is the first study to provide a direct comparison between Bach1 and Bach2, and further to identify expression of both in dopaminergic neurons of the human substantia nigra, an area of the brain. Bach2, in particular, presented as a robust inhibitor of Nrf2-mediated antioxidant responses. Furthermore, Bach2 expression increased with differentiation of model neuroblastoma cells with a dopaminergic phenotype, coinciding with a decrease in full-length Bach1 expression and a blunting of Nrf2 responses. Overall these results support Bach2 as the major inhibitor of Nrf2-mediated antioxidant expression and present Bach2 as a novel target for promoting antioxidant responses and subsequent protection of dopaminergic neurons.

The work described here compared the relative abilities of Bach1 and Bach2 to alter downstream cellular expression of NQO1, HO1, GCLM, GCLC, and GSR as measures of effectiveness. Our results support Bach1 is a mild Nrf2 inhibitor, mostly mediating the expression of HO1, while Bach2 expression has a more profound inhibitory effect. Previous work supports Bach1 as the major mediator of HO1 expression (Kitamuro et al., 2003; Sun et al., 2002; Suzuki et al., 2003), and some studies suggest that Bach1 can modulate GCLM, in addition to HO1 (Warnatz et al., 2011). Differences in methods used may explain the difference in response, as Warnatz and colleagues identified their responses through RNA analyses (gene chip and quantitative real-time PCR) (Warnatz et al., 2011). We cannot, therefore, rule-out mRNA and protein stability differences that may result in differences from the mRNA profile, or that the RNA analyses provide a more sensitive response that could not be adequately observed with protein expression analysis. Further, all of our studies, except where otherwise indicated, were performed in the presence of 5% O_2_, which can alter antioxidant expression (Chapple et al., 2016). Nevertheless, our results indicate Bach2 expression provides a more profound downstream effect compared to Bach1 in limiting antioxidant responses.

We also identified the CNC-bZIP domain is the main requirement for Bach1 and Bach2 function, which is not surprising given that in the absence of this domain, neither protein would bind to ARE promoter sites. However, swapping the CNC-bZIP domains of Bach1 and Bach2 only partially altered their relative activities, indicating that other factors are also responsible for the differences between Bach1 and Bach2. Some of these differences may be further complicated by alterations in nuclear import and export domains, some of which are located in the CNC-bZIP region of Bach1 (Suzuki et al., 2003). The current study was limited by analysis of downstream antioxidant enzyme expression, rather than identifying protein localization or chromatin binding to ARE regions. Future studies could identify the additional mechanic differences between Bach1 and Bach2 or more specific regions responsible for their differences in activity.

Similar to previous reports (Hoshino & Igarashi, 2002; Shim et al., 2006), we show increased Bach2 expression with neuronal differentiation of neuroblastoma lines. This coincided with a blunting of Nrf2 responses after differentiation. Although mild responses to MMF and CoPP could be observed, the overall affect was modest, especially compared to undifferentiated cells. Of note, Nrf2 expression also increased only mildly in differentiated cells in response to MMF-induced activation, and neither CoPP nor CdCl_2_ effectively increase Nrf2 expression after differentiation. Nrf2 expression is regulated by its interaction with Keap1, which targets Nrf2 for proteasomal degradation; however, with Nrf2 activation, this interaction is inhibited, resulting in increased Nrf2 expression and activity (Kobayashi, Kang, et al., 2004; Kobayashi, Ohta, & Yamamoto, 2004; Motohashi & Yamamoto, 2004; Uruno & Motohashi, 2011).

We therefore, cannot rule out a loss of the effectiveness of Nrf2 activators to obstruct the Nrf2-Keap1 interaction in differentiated neurons as a contributing factor to the observed downstream antioxidant enzyme and Nrf2 expression. Further studies using siRNA or genomic knockdown/ knockout of Bach2 would provide additional insights into the direct function of Bach2 in dopaminergic neurons.

The increase in Bach2 after differentiation coincided with a decrease in full-length Bach1 and an increase in a truncated or splice-variant form of Bach1. A splice variant of Bach1 had been identified to omit the DNA-binding domain and the CLS region, but maintain the N-terminus, functioning to help transport full-length Bach1 protein into the nucleus (Kanezaki et al., 2001). The Bach1 lower molecular weight forms in neuroblastoma cells may therefore be this splice variant or function similarly. The current study showed promiscuity between the Bach1 and Bach2 BTB domains. This result suggests that a truncated Bach1 may also transport Bach2, and they may function in tandem to promote their transcriptional repressor function. Since HEK293 cells express Bach1 under basal conditions, endogenous Bach1 may interact with transfected Bach2 to produce the responses we observed. While additional characterization studies are required to fully understand the activities of Bach1 and Bach2 in dopaminergic neurons, the present study identifies both Bach1 and Bach2 as potentially effective transcriptional inhibitors, working in cooperation at mediating Nrf2’s protective effects.

Recent work has supported the roles of antioxidant enzymes and Nrf2 activators as potential therapeutics for the treatment of Parkinson’s disease (Ahuja et al., 2016; Gazaryan & Thomas, 2016; Kaidery et al., 2013; Yang et al., 2009). However, in models of PD, the protective effects are conferred by neighboring glia (Chen et al., 2009; Gan, Vargas, Johnson, & Johnson, 2012; Rojo et al., 2010). Nrf2 was previously identified in the nucleus of substantia nigra neurons in PD brains (Ramsey et al., 2007). The authors posed as one explanation of their results that the presence of Nrf2 may not be potent enough to provide protection to dopaminergic neurons under the stress of neurodegenerative insults. The current work suggests a mechanism by which Nrf2 responses are blunted in dopaminergic neurons, such that Bach2’s presence, in addition to Bach1 or truncated Bach1, may prevent Nrf2’s protective effects in the PD brain. Although our confocal microscopy analyses identified Bach2 as mostly expressed in the soma of the dopaminergic neurons, and no difference in immunoreactivity levels in PD patients versus controls, this result may be due to the analysis of tissue from PD patients who had already experienced severe neurodegeneration and may not be representative of patients in an earlier in a disease state. Our analysis was further limited due to availability of samples. Future studies on additional samples would be necessary to determine if any fine differences were present between the groups and/or if gender or level of PD progression alters Bach2 levels in dopaminergic neurons.

Bach2 has been mostly characterized in B-cell and T-cell lymphocytes (Muto et al., 1998; Muto et al., 2004; Roychoudhuri et al., 2013). Interestingly, B-cells and T-cells express tyrosine hydroxylase and dopamine receptors (Bergquist, Tarkowski, Ekman, & Ewing, 1994; Contreras et al., 2016; Ferrari et al., 2004; Kustrimovic, Rasini, Legnaro, Marino, & Cosentino, 2014; McKenna et al., 2002). Bach2 expression in dopaminergic neurons may therefore be a parallel expression pattern to that of lymphocytes. While Bach2 may not be an initiating cause of Parkinson’s disease, Bach2 likely confers susceptibility to this neuronal population, since they would not likely produce adequate antioxidant enzymes to protect against toxicity and stress. Of note, immunostaining of the cingulate cortex with anti-Bach2 antibodies only yielded non-specific, fluorescent staining that was not detectable at the same exposure as the images captured of the substantia nigra dopaminergic neurons in Figure 3 (data not shown). While a comprehensive analysis of brain regions was not performed, expression of Bach2 may be limited to isolated neuronal populations in the human adult brain.

Overall, the current study provides an initial analysis into the control of antioxidant enzymes in dopaminergic neurons and supports a mechanism whereas Bach2 acts as a robust inhibitor preventing the protective Nrf2-mediated responses. Furthermore, this work shows fundamental differences in response from undifferentiated to differentiated states in neuroblastoma lines, supporting that studies in immortalized cell lines would not be transferable to neurons. In addition, toxicities associated with PD can promote Nrf2 protective responses without a separate inducer (Jakel, Townsend, Kraft, & Johnson, 2007; Yamamoto et al., 2007), suggesting that Nrf2 may have the ability to be active under PD-associated conditions if it were not for transcriptional repressors preventing its function. While further increasing Nrf2 activity through inducers is a potential approach against toxic insults, they would produce blanket increases in antioxidant enzyme expression in multiple cell and tissue populations, while minimally providing protection in the key dopaminergic neuronal population. Our findings provide an alternative approach, whereby chemical inhibitors directed against Bach2 could provide sufficient protective responses to prevent PD progressive neurodegeneration.

## Acknowledgements

This work was supported by Medical Diagnostic Laboratories, LLC, a subsidiary of the Genesis Biotechnology Group, and the Michael J. Fox Foundation for Parkinson’s Disease (E.A.W.). Human brain tissue samples were provided by the Harvard Brain Tissue Resource Center, which is supported in part by PHS grant number R 24 MH 068855. Confocal microscopy was performed with the support and facilities of the University of Pennsylvania Cell and Developmental Biology Microscopy Core.

## Other Acknowledgements

### Author Contributions

All authors had full access to the data in the study and take responsibility for the integrity of the data and the accuracy of the data analyses. *Conceptualization, Methodology, Validation, Investigation, Formal Analysis, Resources, Data curation, Writing, Visualization, Supervision, Project Administration, and Funding acquisition*: E.A.W.

## Graphical Abstract

Transcriptional repressors Bach1 and Bach2 are both expressed in dopaminergic neurons, inhibiting Nrf2 responses and antioxidant enzyme expression. We found that Bach1 is a modest inhibitor of Nrf2 responses, and Bach2 is a robust inhibitor, suggesting that Bach2 presence in dopaminergic neurons may prevent expression of protective antioxidant enzymes and contribute to progressive neurodegeneration.

